# A Continuum Fibre Network Model for the Anisotropic and Compressible Mechanics of Human Skin

**DOI:** 10.64898/2026.11.06.764854

**Authors:** Conall Quinn, Mahtab Vafaeefar, Behrooz Fereidoonnezhad, Ted J. Vaughan

## Abstract

We present a Continuum Fibre Network (CFN) formulation for modelling anisotropy, strain stiffening and volume change in fibrous soft tissues. The model combines a three-dimensional fibre orientation distribution, a tension-only fibre law and a separate phenomenological volumetric potential. We calibrated the model using published uniaxial tensile and volume-change data for excised human skin from separate studies. Changes in cross-sectional area affected the tensile stress estimates for models that predicted volume change. Simulations of indentation, shear and biaxial tension illustrate predictions of the calibrated formulation under additional loading conditions. The CFN formulation included a weak isotropic neo-Hookean ground matrix with a prescribed shear modulus of 0.1 MPa.

## 1 Introduction

Skin serves as a protective barrier for the body, shielding underlying tissues from mechanical, chemical, biological, and thermal influences [1]. Its structure varies significantly depending on anatomical location, age, gender, race, and health [2-6]. Skin consists of three layers: the epidermis, dermis, and hypodermis[7]. The epidermis is the outermost layer (75-150μm thick) that prevents water loss and protects against environmental damage and pathogens [8]. The dermis is the middle layer (1.3 – 2.4mm thick), composed of collagen and elastin fibres, and provides structural support and elasticity. The hypodermis, the innermost layer, acts as a shock absorber and anchors the skin to underlying structures [9, 10]. Skin consists of an extracellular matrix (ECM) and fluid components. The ECM is comprised of a network of collagen and elastin fibres along with proteoglycans that provide mechanical support. Fluid components facilitate nutrient transport and distribute mechanical loads. Elastic fibres enable reversible deformation at low strains [11] while collagen fibres maintain tissue integrity at high strains. Proteoglycans consist of a core protein with glycosaminoglycan (GAG) chains forming the ground matrix that retains water and enhances resistance to compressive forces [12]. This hierarchical, fibrous, and fluid-saturated architecture governs the nonlinear, hyperelastic, anisotropic, and compressible mechanical response of skin, presenting significant challenges for constitutive modelling [13].

Accurately capturing this behaviour in finite element analyses requires constitutive models capable of representing the underlying microstructural mechanisms of soft tissues, with various examples across different tissues such as blood vessels [14, 15], heart tissue [16, 17], ligaments [18, 19] and skin itself [20, 21]. Hyperelasticity in skin arises from the initial waviness of collagen fibres, their progressive reorientation, and fully straightening under tension [22, 23], while anisotropy arises from preferential fibre alignments (Langer lines), varying by anatomical location [24-27]. Volume changes in skin can reflect the combined behaviour of the solid structure and interstitial fluid [28-30]. Traditional hyperelastic models developed [31-33] for rubber-like materials tend to oversimplify key features of biological tissues, including fibre mechanics, anisotropy, and porosity. To address these limitations, fibre-reinforced constitutive models often employ a single continuous strain energy function using exponential or power-law terms to approximate the collagen fibres’ characteristic J-shaped stress-strain response [13, 33, 34]. Gasser et al. [35] proposed a continuum model for arterial tissue with separate ground matrix and fibre contributions. These models assume perfect bonding and stress equilibrium between fibres and matrix, simplifying their interaction. Although the model can reproduce experimental responses under uniaxial loading, the predicted transverse and volumetric responses also depend on the selected constitutive terms. A tensile fit alone does not establish predictive accuracy beyond its calibration range. The chosen fibre-law form controls the rate of stiffening and must be assessed over the relevant strain range. Two-family and distributed-orientation formulations impose different architectural assumptions. Restricting fibre orientations to two dimensions ignores out-of-plane contributions and changes the matrix contribution to transverse load balance [20]. An incompressibility constraint excludes the volume changes reported in experiments [36-39]. More realistic volumetric functions such as those by Moerman et al. [40] better represent the transition from compressible to nearly incompressible states in soft tissues. Fibre–matrix stress equilibrium can contribute to transverse expansion under tension. Fereidoonnezhad et al. [41] demonstrated that excessive matrix compression along one axis leads to unrealistic out-of-plane expansion driven by stress equilibrium. The strong sensitivity to certain model parameters warrants careful assessment and motivates constitutive frameworks that more directly reflect underlying fibre network mechanics.

Recent advances in constitutive modelling have sought to improve soft tissue mechanics representation and eliminate unphysical behaviours. Nolan et al. [42] modified the fibre strain invariant in the Holzapfel-Ogden-Gasser (HGO) model, enabling fibres to carry volumetric loads. The Gasser-Ogden-Holzapfel (GOH) model further enhanced anisotropic behaviour by incorporating three-dimensional fibre dispersion [35]. While this statistical approach effectively captures macroscopic anisotropy, it averages fibre orientations through structure tensors rather than integrating each reference direction individually. Fereidoonnezhad et al. [41] introduced a bilinear strain-stiffening hyperelastic model providing greater control over fibre and matrix mechanical responses. Despite its flexibility, minimising auxetic effects demands precise matching of fibre and matrix stiffness, complicating parameter identification. Discrete fibre network models explicitly simulate individual fibre interactions and capture complex microstructural phenomena including nonlinear deformation, fibre recruitment, frictional contact, network densification, and damage evolution [43]. Their computational cost restricts their use in large-scale applications. These approaches differ in their resolution of fibre interactions and computational requirements.

The objective of this study was to develop a continuum fibre network (CFN) formulation with separate control of fibre orientation, strain stiffening and volumetric response. The formulation builds on existing approaches in which fibre stretch depends on the full deformation [42], the fibre response follows a tension-only stiffening law based on a bilinear formulation [41], and the volumetric response follows a phenomenological law [40]. Calibration used published tensile and volume-change measurements; additional loading simulations assessed model responses rather than independent experimental validation. Fibre reinforcement was represented by continuum orientation averaging without explicitly resolving network connectivity.

## 2 Methodology

### 2.1 General Framework for Fibre-Reinforced Soft Tissues

Soft tissue deformation is characterised by the deformation gradient **F**, with the right Cauchy– Green tensor defined as:

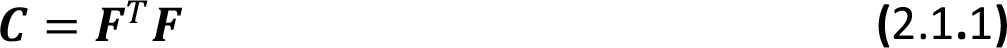

Volumetric change is described by the Jacobian:

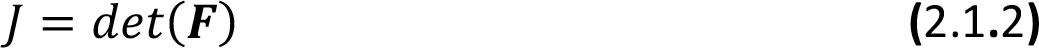

which gives the local ratio of deformed to reference volume. In general fibre-reinforced constitutive models, the strain energy (*Ψ*) is expressed as the sum of an isotropic matrix contribution (*Ψ*_*m*_) and an anisotropic fibre contribution (*Ψ*_*fi*_):

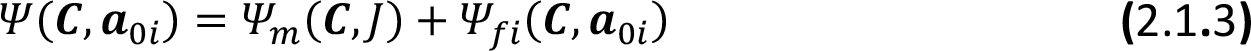

The matrix phase typically governs bulk and distortional response, often enforcing near-incompressibility through a volumetric penalty function dependent on *J*. The fibre phase introduces directional stiffness through invariants of the form:

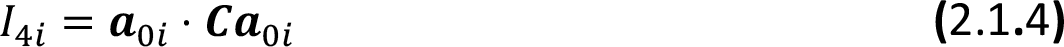

where ***a***_0 *i*_ denotes the reference fibre orientation. Fibres are generally assumed to carry load only in tension, contributing anisotropic stress that is superimposed onto the isotropic matrix stress. This matrix-fibre separation enables flexible fitting of nonlinear and anisotropic behaviour, but it also imposes structural constraints. Phenomenological bulk terms can describe volumetric resistance at the continuum scale. Furthermore, enforcing stress equilibrium between a compliant matrix and stiff fibres can introduce artificial transverse expansion or auxetic effects under uniaxial tension. These limitations motivated a CFN formulation with separate descriptions of fibre orientation and stiffening, isotropic matrix resistance, and resistance to volume change.

### 2.2 Continuum Fibre Network (CFN) Model

The CFN energy per unit reference volume adds an isochoric matrix potential, orientation-weighted fibre energies and a volumetric potential.

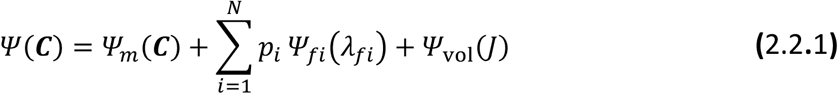

The unweighted energy of fibre direction *i* is *Ψ*_*fi*_(*λ*_*fi*_). Its reference orientation probability *p*_*i*_ enters the sum once. The separate potential *Ψ*_vol_(*J*) describes volumetric resistance. Reference directions were sampled on a Fibonacci sphere using the polar angle *ϕ*_*i*_ and the azimuth *θ*_*i*_.

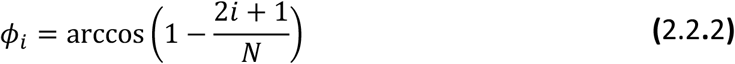

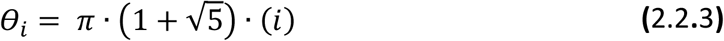

where *i* = 0, 1, 2, …, *N* − 1 and *N* is the total number of fibres. Spherical coordinates were mapped to three-dimensional Cartesian space using:

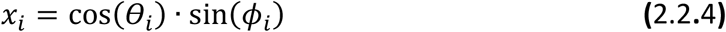

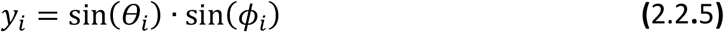

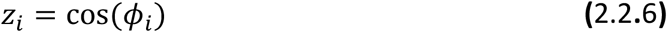

Each fibre position satisfies the unit sphere constraint:

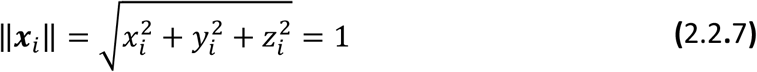

Reference fibre directions were sampled quasi-uniformly on the unit sphere using a Fibonacci lattice. A generalised ellipsoidal exponential weighting assigned the reference orientation probabilities. Its shape was described by:

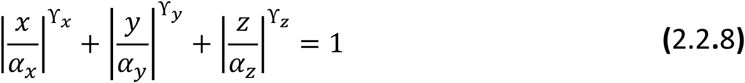

The parameters *α*_*x*_, *α*_*y*_, and *α*_*z*_ controlled the spread of fibre orientations along the three principal axes, while ϒ_*x*_, ϒ_*y*_, ϒ_*z*_ governed the directional decay of the weighting function. For a quadratic exponent in the fibre-direction components, the standard Bingham weighting was:

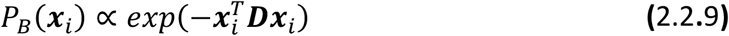

The anisotropic scaling matrix ***D*** controlled the directional bias:

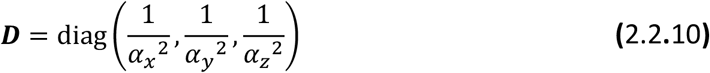

Allowing directional exponents gave the generalised ellipsoidal weighting:

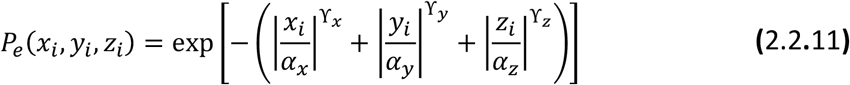

The exponential weighting favoured preferred fibre orientations. The directional exponents ϒ_*x*_, ϒ_*y*_, ϒ_*z*_ controlled the weighting shape. When all three exponents satisfied ϒ = 2, the weighting had a quadratic Bingham form. The sampled weights were normalised so that their probabilities summed to one:

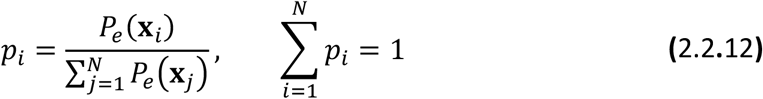

To capture preferential in-plane alignment, a rotation transformation was applied, reorienting fibre directions within the x-y plane:

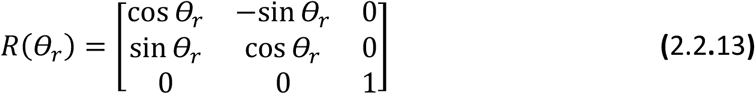

where *θ*_*r*_ represents the preferred fibre alignment relative to the x-axis. The transformed fibre position is given by:

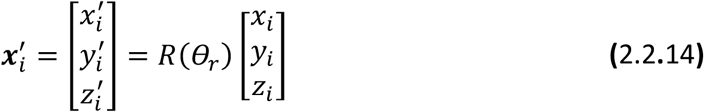

The reference weights were evaluated and normalised before rotation. Each weight remained attached to its fibre direction when the network was rotated. Dispersion was quantified using the probability-weighted variance of the fibre-direction components along each principal axis:

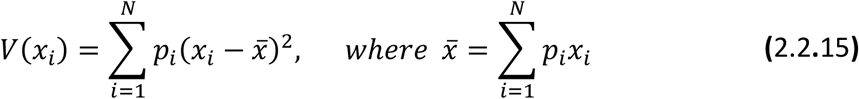

Repeating this process for *y*_*i*_ and *z*_*i*_ yielded the corresponding variances *V*(*y*_*i*_) and *V*(*z*_*i*_), defining the directional anisotropy of the fibre network. **Figure 1** illustrates the direction sampling and rotation.

**Figure 1.**
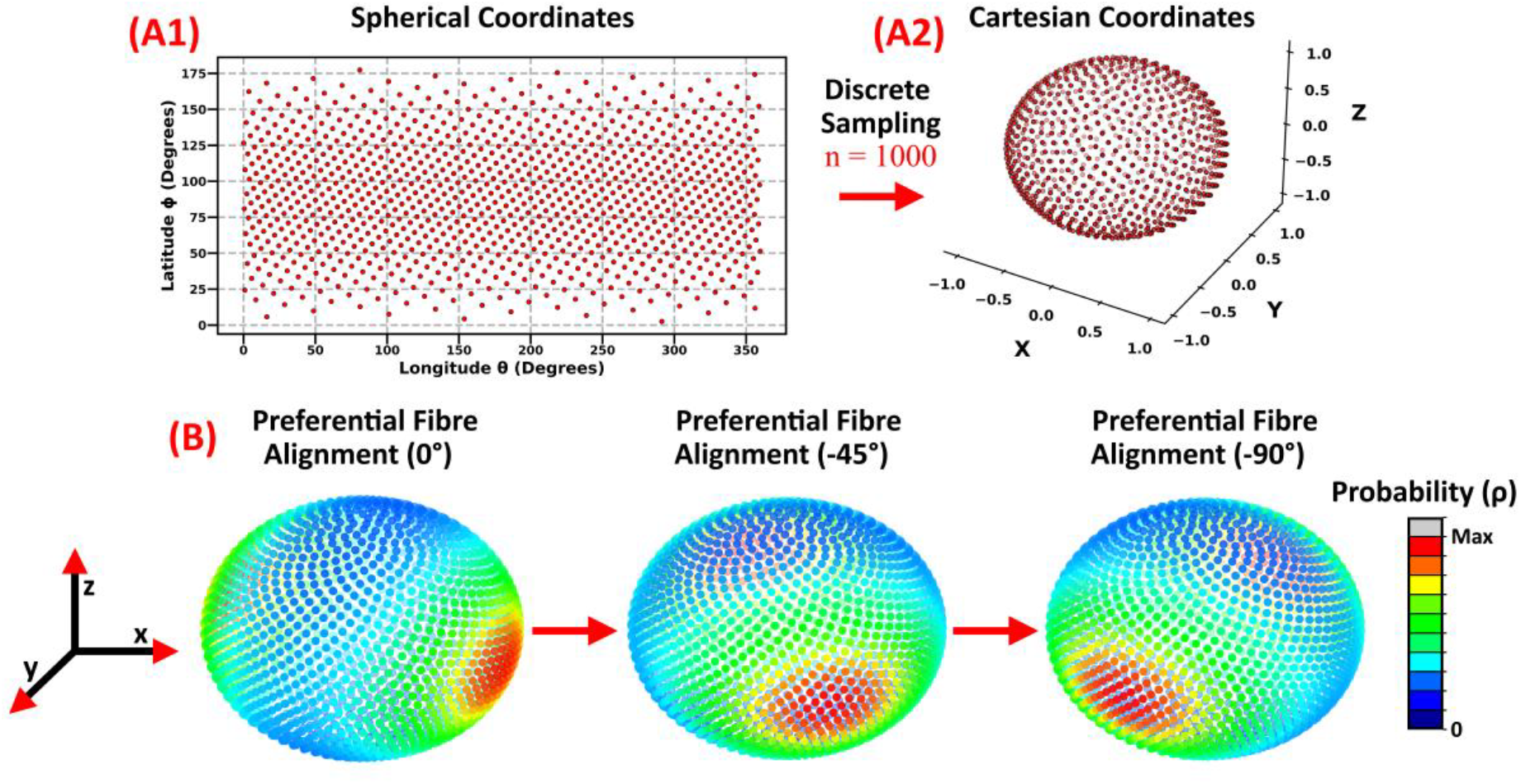
**(A1)** Distribution of 1000 fibre orientations in spherical coordinates. **(A2)** Mapping to Cartesian coordinates forms a quasi-uniform unit sphere distribution. **(B)** Fibre alignment adjusted by rotation (0^°^, −45^°^, −90^°^), with probability density visualised (red = high, blue = low). The illustrative anisotropic ellipsoidal Bingham (AEB) distribution uses semi-axes *α*_*x*_ = 2, *α*_*y*_ = 0.81, and *α*_*z*_ = 0.7357, with ϒ_*x*_ = ϒ_*y*_ = ϒ_*z*_ = 2.

The CFN uses the full fibre invariant *I*_4*i*_ = ***a***_0*i*_ ⋅ ***Ca***_0*i*_ and the stretch 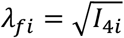. This allows fibre stretch to depend on volume change. The isochoric matrix term supplies distortional resistance.

Fibre mechanics were described by a tension-only strain-stiffening law following Fereidoonnezhad et al. [41]. The scalar fibre coefficient *g*_*i*_ comprised an initial linear region, a quadratic transition and a final linear region. The fibre-law coefficient was set to zero at or below unit fibre stretch. The tensile response was:

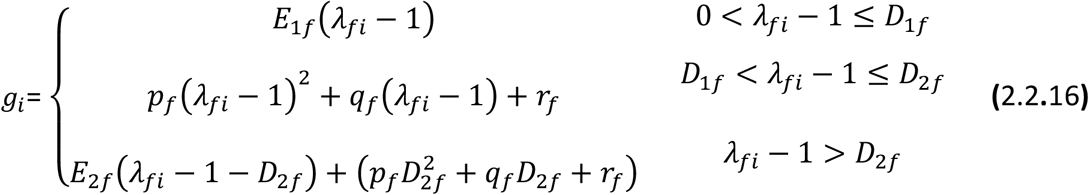

Here *E*_1*f*_ and *E*_2*f*_ controlled the initial and final slopes, while *D*_1*f*_ and *D*_2*f*_ were positive transition strains (see **Figure 2(A2)). The coefficients** *p*_*f*_, *q*_*f*_ and *r*_*f*_ matched the coefficient and its first derivative at both positive-strain joins. These interpolation coefficients were distinct from the orientation probabilities:

**Figure 2.**
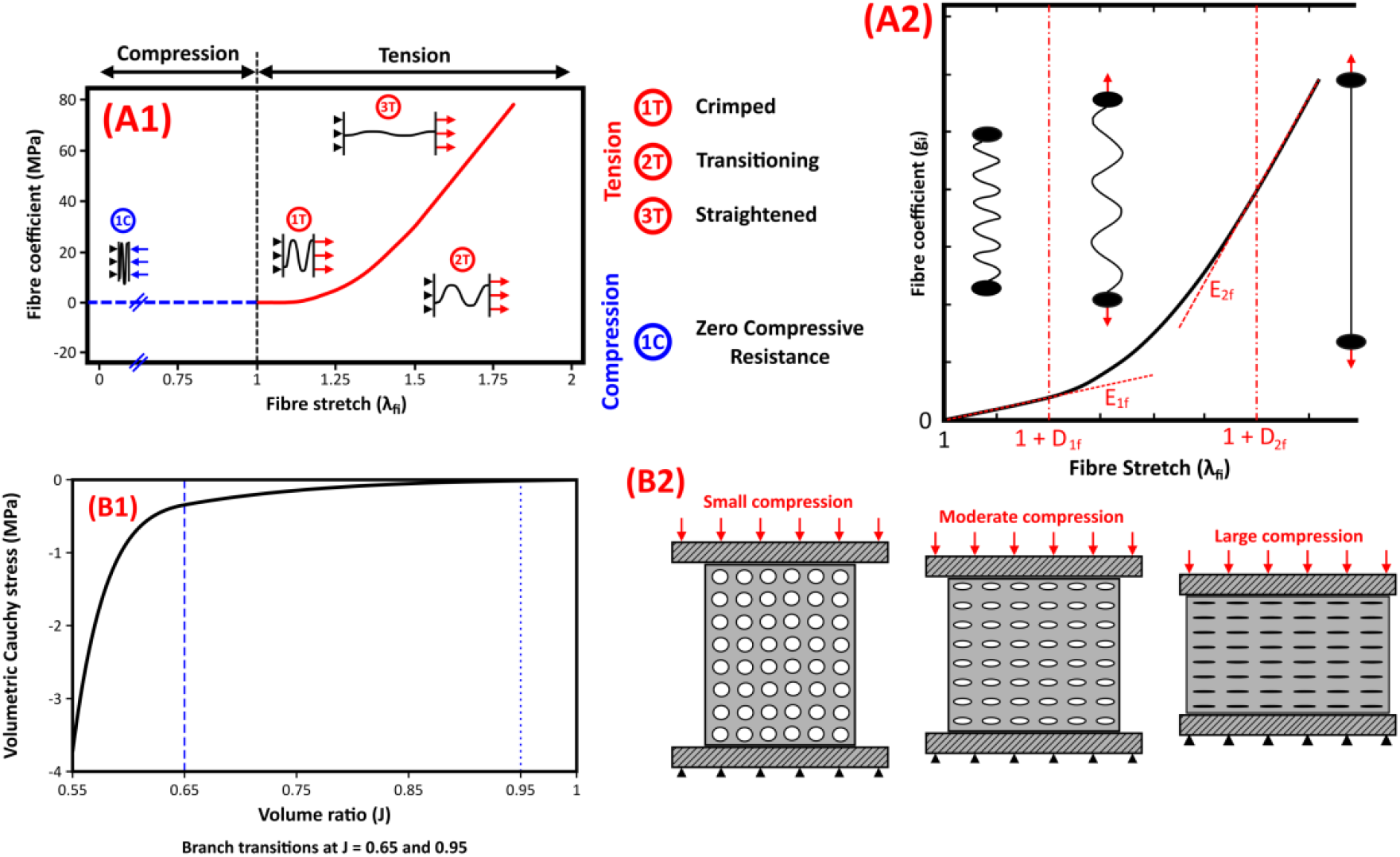
**(A1)** Schematic of the assumed fibre response in tension and compression. The fibre contribution is active only above unit fibre stretch. **(A2)** Tension-only fibre coefficient with two linear branches, a quadratic transition and parameters *E*_1*f*_, *E*_2*f*_, *D*_1*f*_, *D*_2*f*_. (**B1**) Volumetric Cauchy stress from the prescribed compression law (*J* < 1). Dashed guides mark the branch transitions. (B2) Schematic illustration of decreasing volume.

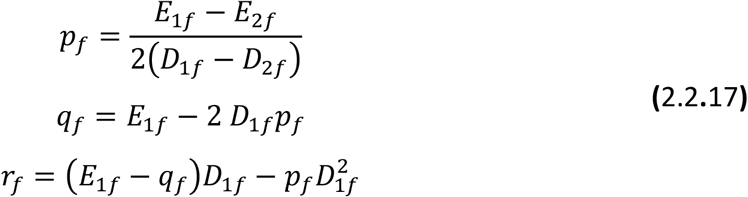

A fibre contributes only above unit fibre stretch. For each fixed reference direction, the affine current vector was *a*_*i*_ = *F a*_0*i*_, and its norm defined the fibre stretch. The orientation-weighted fibre sum formed a Kirchhoff stress; division by the volume ratio gave the fibre-network Cauchy stress (*σ*_*network*_):

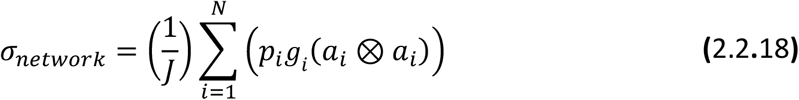

This formulation represents distributed directional reinforcement and strain stiffening without resolving network connectivity. Near-incompressible formulations restrict volume change through a bulk modulus *κ*. However, the porous architecture of biological tissues leads to measurable volumetric changes during mechanical loading, as reported across different soft tissues including myocardium and skin [28, 44, 45], highlighting the need for a more detailed volumetric representation.

The CFN volumetric response was prescribed separately for compression (*J* < 1) and expansion (*J* > 1):

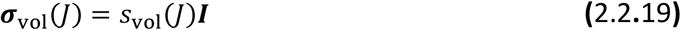

The signed scalar *s*_vol_(*J*) is negative in compression, positive in expansion, and zero at *J* = 1. Appendix A defines the two branch laws. The fitted compression law has its first threshold at *J* = 1, so its initial linear interval reduces to the reference state. Appendix F integrates the signed stress to obtain the volumetric energy. The law describes resistance to volume change phenomenologically.

The CFN formulation included an isotropic neo-Hookean matrix component with shear modulus *μ* = 0.1 MPa:

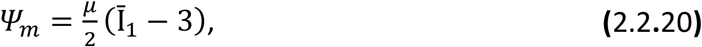

where 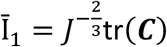. The matrix provided distortional resistance when fibres were inactive, while the volumetric potential resisted volume change. The total stress was derived from the combined matrix, fibre and volumetric energies (Appendix F). The combined Cauchy stress is given in Equation **(F.5)**.

### 2.3 Numerical Implementation

The CFN constitutive formulation was implemented as a user-defined material (UMAT) subroutine within Abaqus (Dassault Systèmes Simulia Corp. [46]). Anisotropic ellipsoidal Bingham (AEB) fibre orientations and associated probability weights were generated using a custom Python script and exported as a text file for import via the UEXTERNALDB subroutine. This procedure enabled consistent integration of statistically defined fibre architectures within the finite element framework. Initial statistical analysis and visualisation of fibre distributions, as well as inspection of the nonlinear volumetric scaling functions, were performed in Python to support model verification and interpretation. Material parameter identification was performed using an automated Python-Abaqus workflow. The earlier tensile calibration used for the comparison cases followed these steps:

1. Update UMAT parameters
2. Execute simulations
3. Compute the coefficient of determination *R*^2^ between simulated and experimental responses
4. Optimise material parameters using the Limited-memory Broyden–Fletcher– Goldfarb–Shanno with Box constraints (L-BFGS-B) algorithm

The earlier tensile calibration used the squared stress error:

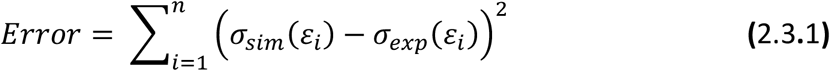

where *σ*_*sim*_ and *σ*_*exp*_ denote simulated and experimental stresses at strain *ε*_*i*_.

### 2.4 Baseline Material Calibration

One set of CFN coefficients was calibrated to the parallel and perpendicular tensile curves and the mean volume-change curve, evaluated against the parallel simulation. The tensile and volume targets came from separate studies. Simulated responses were compared at measured gauge stretch.

#### 2.4.1 Fibre Mechanical Parameter Identification

The human back-skin tensile tests used dogbone specimens prepared to ASTM D412 [47]. The parallel and perpendicular tensile targets were digitised from the experimental curves in Figure 10 of Ní Annaidh et al. [20]. Simulated nominal stress was calculated by dividing axial force by the initial cross-sectional area. Tensile fit scores were calculated against the published values as plotted.

The same fibre distribution and constitutive coefficients were used for both dogbone orientations, with the reference directions rotated by 90° for perpendicular loading. A bounded least-squares update varied two ellipsoid semi-axes, the straightening modulus and the fibre strain at the end of recruitment. The objective used equally weighted mean square errors after normalising each target by its measured span, with penalties for endpoint errors and parameter changes.

#### 2.4.2 Fibre Orientation Parameter Identification

The joint calibration adjusted the two transverse semi-axes of the ellipsoidal orientation distribution, which was sampled with 200 directions. The ratio of out-of-plane to in-plane weighted directional variances was defined as:

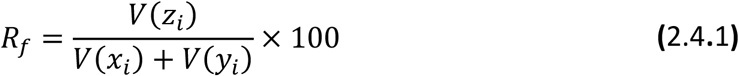

where *V*(*x*_*i*_), *V*(*y*_*i*_), and *V*(*z*_*i*_) denote the calculated weighted directional variances along the x, y, and z axes, respectively.

#### 2.4.3 Volumetric Parameter Calibration

Volumetric parameters were calibrated using experimental measurements reported by Wahlsten et al. [28], who quantified the volumetric compressibility of human skin under uniaxial tension. Their study demonstrated substantial reductions in volume ratio *J* with increasing axial stretch, without uniquely identifying the underlying structural or fluid mechanism.

At 20% axial elongation the digitised mean volume ratio was 0.697. The previously calibrated compression coefficients were fixed during the joint fibre calibration, and the parallel volume curve was included in the objective (see **Appendix A**). The predicted volume response beyond the measured stretch range was treated as extrapolation.

### 2.5 Sensitivity and Directional-Sampling Analysis

A unit cube subjected to uniaxial tension across multiple stretch levels was used to evaluate the influence of collagen fibre mechanics and network architecture on tissue behaviour.

Fibre mechanical parameters were varied systematically, including uncoiling stiffness (*E*_1*f*_), first transition strain (*D*_1*f*_), straightening stiffness (*E*_2*f*_), and second transition strain (*D*_2*f*_). The influence of fibre orientation was assessed by varying preferential alignment (*α*_*x*_) from highly anisotropic configurations (*α*_*x*_ ≫ (*α*_*y*_, *α*_*z*_)) to isotropic distributions (*α*_*x*_ = *α*_*y*_ = *α*_*z*_). The parameters of the volumetric compression law were also varied to assess sensitivity to the prescribed volumetric resistance. Stress anisotropy under isotropic volumetric loading was quantified using the inverse triaxiality ratio:

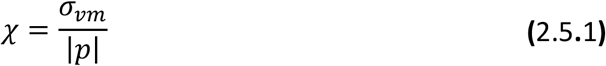

where *σ*_*vm*_ is the von Mises stress and *p* is the hydrostatic pressure. The denominator uses its magnitude. For an isotropic material under pure volumetric deformation, *σ*_*vm*_ = 0 and *χ* = 0 when the pressure magnitude is nonzero. The benchmark assessed residual directional bias from finite fibre sampling. Fibre networks containing 20, 100, 500, 1000, and 10,000 fibres were evaluated to assess isotropy and computational efficiency, with CPU time recorded for each simulation.

### 2.6 Comparative Model Evaluation

The CFN model was compared against BLM, GOH, and HGO fibre-reinforced constitutive models. The reported calibrations used the selected tensile data. The comparison is specific to the reported parameter sets. Equivalent optimisation and parameter identifiability were not established.

The directional stress response was evaluated by subjecting a cube to uniaxial tension while rotating the preferential fibre direction from 0^°^ to 90^°^ in 10^°^ increments within the x–y plane. At axial stretch 1.50, the von Mises stress for each model was normalised by its peak over the angle sweep. Auxetic behaviour and volumetric response were investigated by tracking transverse stretches (*λ*_2_, *λ*_3_) and volume ratio as a function of axial stretch (*λ*_1_). The cube response was evaluated separately from the integrated dogbone gauge volume. The section-average axial stress was calculated by dividing the axial force by the instantaneous cross-sectional area:

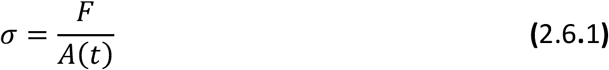

For aligned stretches and near incompressibility (*J* ≈ 1), the current area can be estimated from the exact incompressible relation:

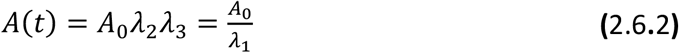

For the direct area calculation, the deformed gauge cross-section was split into two triangles. The edge vectors (**v**_*i*_) were formed from the four deformed nodal positions with a common origin at the first node. The second vector is the shared diagonal, listed twice in Appendix D. The two triangle areas were summed:

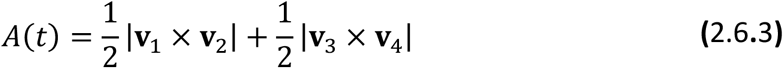

Appendix D gives the nodal construction. The area and edge vectors are given in Equations **(D.1)** and **(D.2)**.

### 2.7 Multi-Axial Loading Analysis

Indentation was simulated using a cylindrical specimen (radius 10 mm, height 10 mm) compressed by a rigid indenter (radius 2.5 mm) to a target displacement of 5 mm. The updated CFN simple-shear simulation imposed 5 mm of horizontal displacement on a 10 mm × 10 mm × 2.25 mm specimen. The biaxial tests used a cruciform specimen with a 10 mm × 10 mm central region and opposing grips displaced by 2.5 mm per edge along each axis.

## 3 Results

### 3.1 CFN Calibration and Parameterisation

**Table 3.1.1** lists the CFN parameters used in **Figure 3**. Histological observations informed the initial orientation distribution [20]. The parallel and perpendicular tensile fits gave RMSEs of 0.874 MPa and 1.040 MPa (R^2^ = 0.996 and 0.993, respectively). The fit to the mean volume-change curve from Wahlsten et al. [28] gave an RMSE of 0.0140 (R^2^ = 0.980).

**Table 3.1.1.** CFN parameters for the joint tensile and volume calibration. Fibre and volumetric stress coefficients are in MPa.

|  | **α<sub>x</sub> / α<sub>y</sub> / α<sub>z</sub>** | **ϒ<sub>x</sub> / ϒ<sub>y</sub> / ϒ<sub>z</sub>** | **Fibre Number** |  |
|---|---|---|---|---|
| **Fibre Orientation** | 2 / 0.907 / 0.702 | 2 / 2 / 2 | 200 |  |
|  | **D<sub>1f</sub> / D<sub>2f</sub>** | **E<sub>1f</sub> / E<sub>2f</sub>** |  |  |
| **Fibre Mechanical** | 0.0175/ 0.291 | 2 / 549.575 |  |  |
|  | **J<sub>1c</sub> /E<sub>1c</sub>** | **J<sub>2c</sub> /E<sub>2c</sub>** | **J<sub>3c</sub> /α<sub>c</sub>** | **β<sub>c</sub>** |
| **Volumetric (Compression)** | 1 / 0.288 | 0.95 / 0.243 | 0.65 / 3.568 | 25 |
|  | **J<sub>1e</sub> /E<sub>1e</sub>** | **J<sub>2e</sub> /E<sub>2e</sub>** | **J<sub>3e</sub> /α<sub>e</sub>** | **β<sub>e</sub>** |
| **Volumetric (Expansion)** | 1.2 / 0.1 | 1.4 / 0.01 | 8 / 0 | 2 |

**Figure 3** shows the fitted fibre distribution and the tensile and volume responses. Histological observations from Ní Annaidh et al. [20] informed the initial distribution. The 29.8% out-of-plane to in-plane variance ratio in **Figure 3(B2)** was calculated from the modelled fibre directions. Ní Annaidh et al. [20] reported binned histological orientation frequencies. At the largest perpendicular stretch in **Figure 3(C1)**, the CFN predicted 36.86 MPa, compared with 36.92 MPa from the published curve.

**Figure 3.**
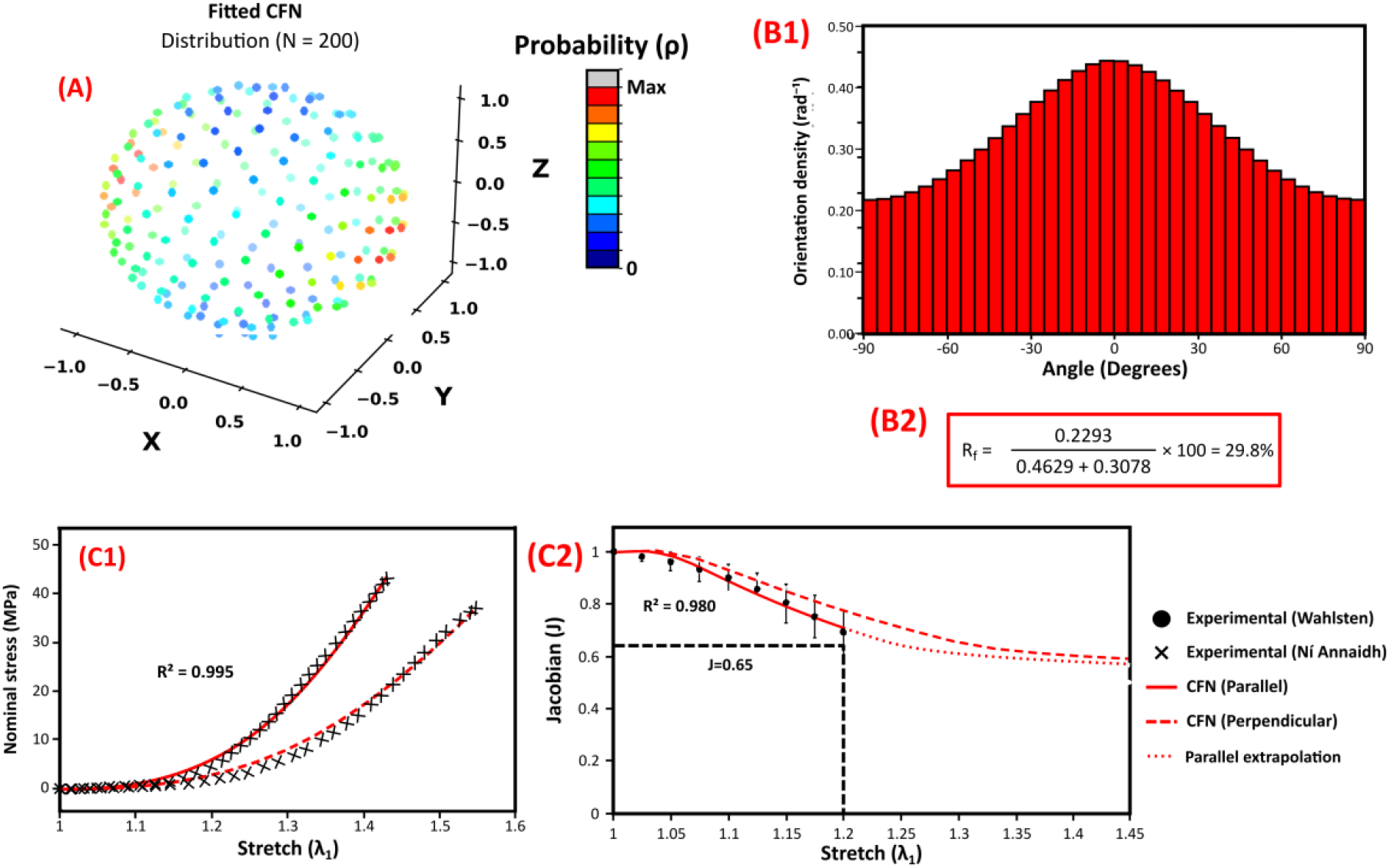
**(A)** Ellipsoidal fibre distribution with 200 directions. **(B1)** In-plane orientation density in 5° bins. **(B2)** Ratio of out-of-plane to in-plane weighted directional variance. **(C1)** Simulated nominal stress–stretch curves for parallel (solid) and perpendicular (dashed) loading with digitised tensile data and a pooled R^2^. **(C2)** Gauge volume ratio and mean volume-change data from Wahlsten et al. The dotted parallel segment extrapolates beyond stretch 1.20. The perpendicular volume curve is a prediction. The horizontal guide marks the terminal compression threshold.

### 3.2 Constitutive Model Comparison

The BLM, GOH and HGO parameters are listed in **Table 3.2.1**. The tensile fits shown in **Figure 4(B–D)** have R^2^ values of 0.994, 0.807 and 0.994, respectively. The runs behind these fits could not be verified. Figures 4–7 show the updated CFN curves. Figures 8 and 9 show the earlier contours.

**Table 3.2.1.** Reported parameter sets for the earlier HGO, GOH and BLM comparisons.

|  | **C<sub>10</sub> (MPa)** | **D<sub>1</sub> (MPa<sup>−1</sup>)** | **k1 (MPa)** | **k2** | **ϕ** | **Fibre Angle (°)** |
|---|---|---|---|---|---|---|
| **HGO** | 1 | 0.919 | 120 | 0 | 0.1 | 40 |
|  | **C<sub>10</sub>** | **D<sub>1</sub>** | **k1** | **k2** | **κ** | **Fibre Angle** |
| **GOH** | 1 | 0.0001 | 620 | 0 | 0.2 | 42 |
|  | **λ<sub>1m</sub> / λ<sub>1f</sub>** | **λ<sub>2m</sub> / λ<sub>2f</sub>** | **E<sub>1m</sub> / E<sub>1f</sub>** | **E<sub>2m</sub> / E<sub>2f</sub>** | **D<sub>1</sub>** | **β** |
| **BLM** | 0.03 / 0.013 | 0.11 / 0.078 | 0.03 / 0.2 | 300 / 230 | 0.919 | 1.1 |

**Figure 4.**
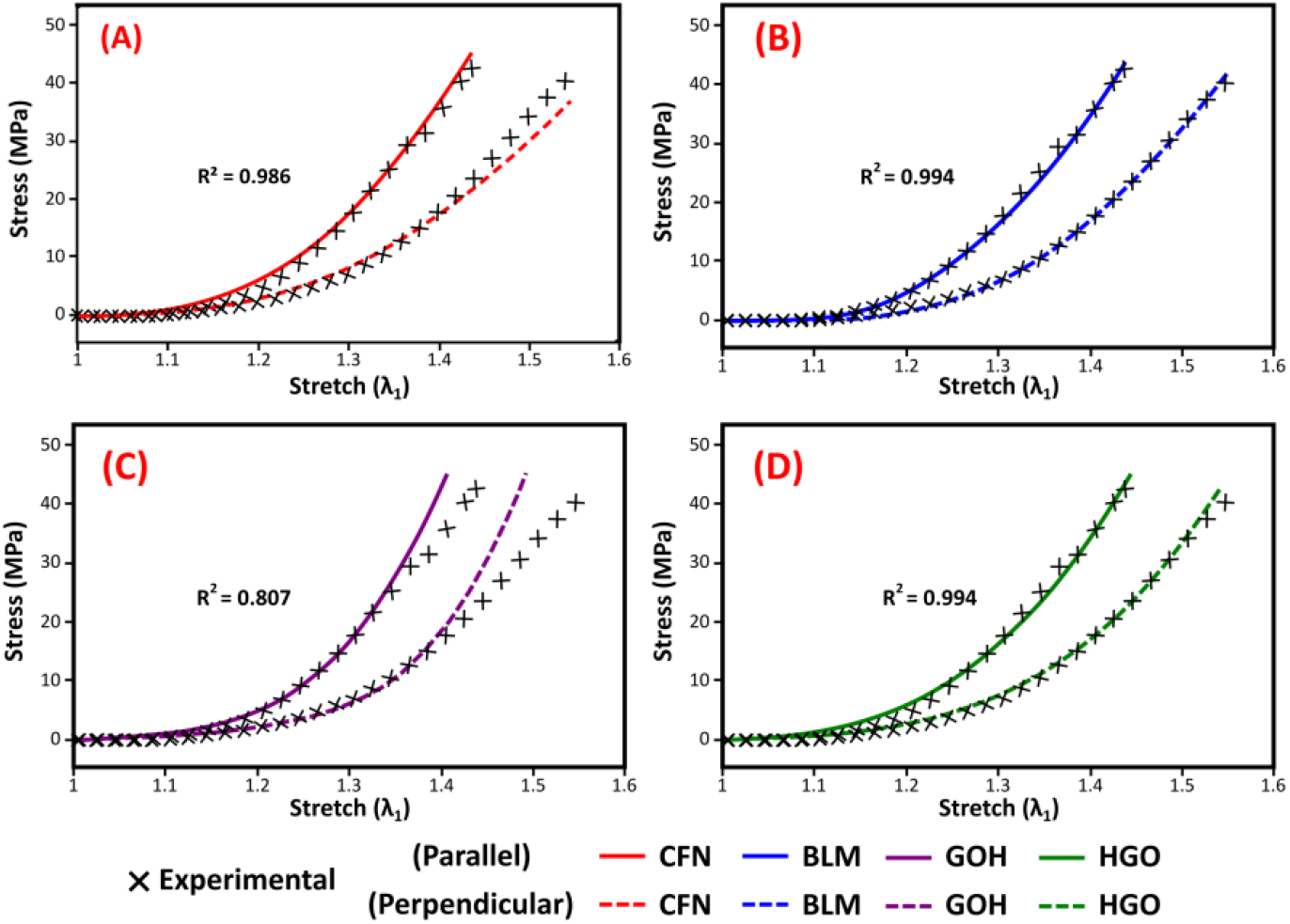
Stress–stretch responses for human skin in parallel and perpendicular tension. **(A)** Updated CFN curves with a pooled R^2^ of 0.986 for the experimental points shown. Figure 3(C1) reports the fit to the calibration data. **(B)** BLM, **(C)** GOH and **(D)** HGO tensile fits from the earlier comparison. Ní Annaidh et al. [20] reported the histological orientation distributions.

**Figure 5** shows the anisotropic mechanical response for the CFN model with a three-dimensional AEB distribution, the BLM model with a two-dimensional von Mises distribution, and the two-fibre GOH and HGO models. The CFN and BLM responses were greatest in the prescribed preferred direction (0°) and decreased towards the perpendicular direction (90°). In contrast, the prescribed fibre angles for the GOH and HGO models were 42° and 40°, respectively (see **Table 3.2.1**). Their displayed stress maxima did not coincide with the prescribed preferred direction (0°).

**Figure 5.**
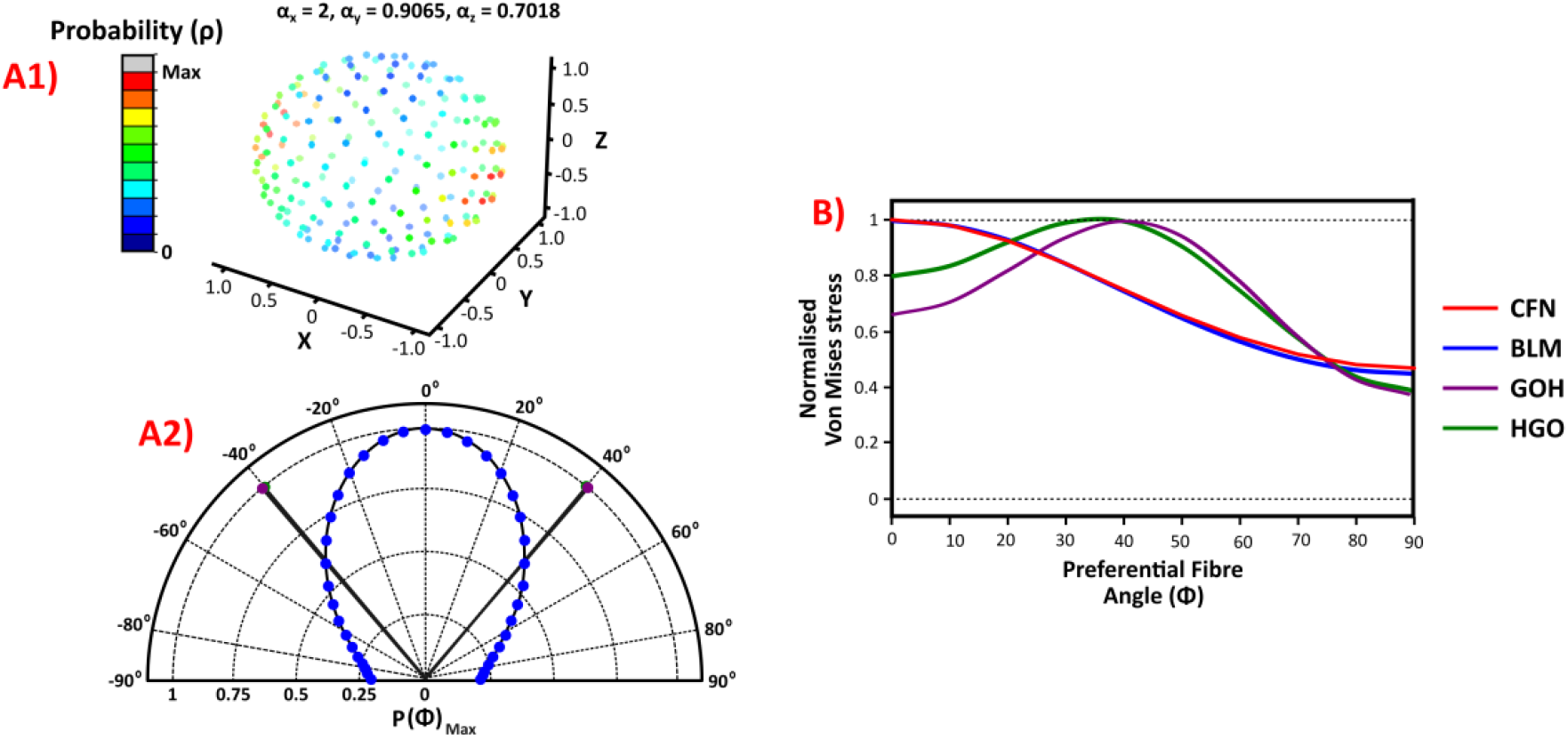
Fibre orientation distributions used for the comparison in **Figure 4**. **(A1)** Three-dimensional CFN distribution. **(A2)** BLM von Mises orientation distribution and GOH (purple, ±42°) and HGO (green, ±40°) fibre angles. **(B)** Cube von Mises stress at axial stretch 1.50 across the 0°–90° orientation sweep, normalised by each model’s own peak. The red curve uses the updated CFN coefficients.

**Figure 6** shows transverse expansion in the selected compressible BLM and HGO cube simulations, with greater out-of-plane deformation (*λ*_3_) compared to the transverse direction (*λ*_2_) under uniaxial stretch. These predictions depend on the selected fibre, matrix and volumetric terms. The CFN model employs a modified version of the piecewise volumetric strain-energy law originally proposed by Fereidoonnezhad et al. [44], introducing additional scaling functions to control volumetric stiffening at large deformations.

**Figure 6.**
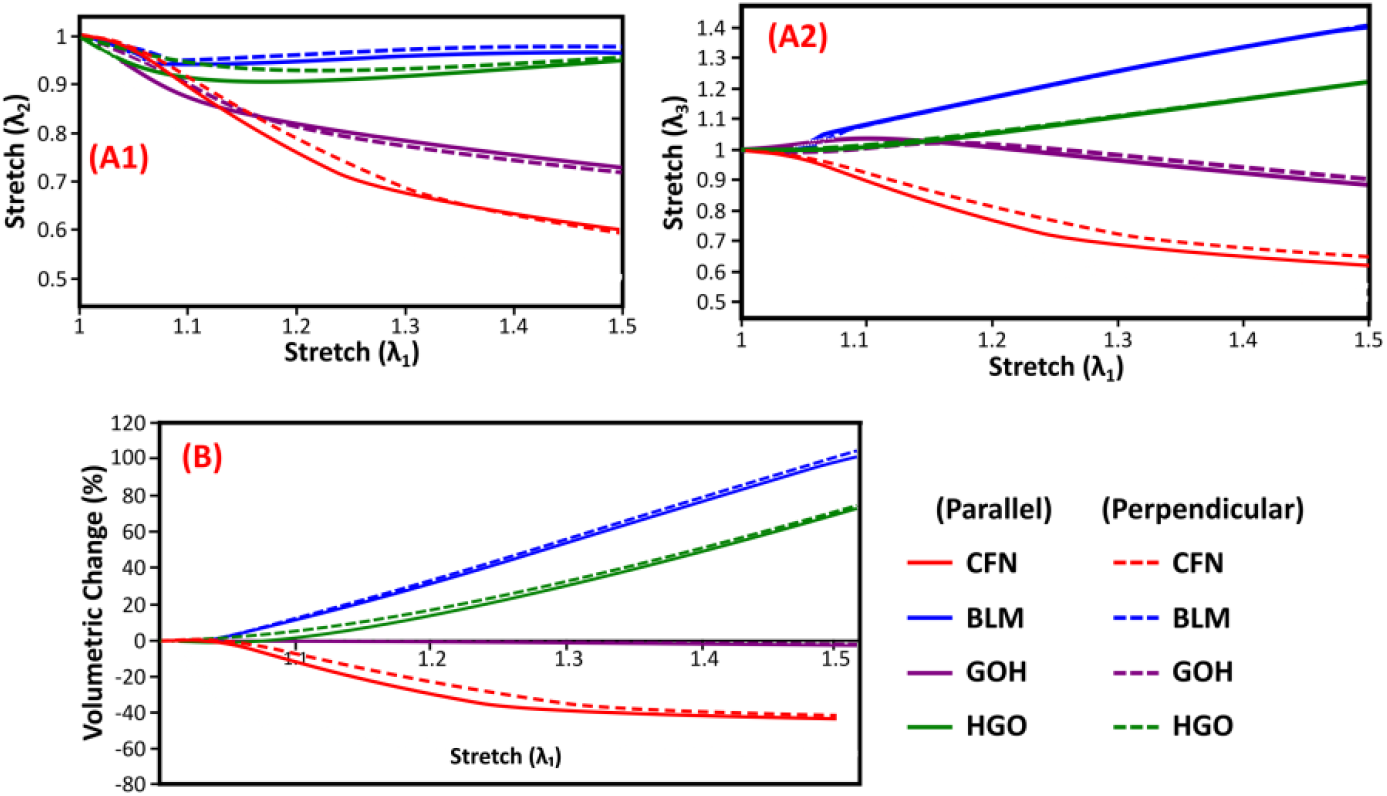
Transverse and volumetric responses of the cubes under uniaxial tension. **(A1)** Width stretch. **(A2)** Thickness stretch. **(B)** Volume change. The CFN curves use the joint calibration; BLM, GOH and HGO are archived comparisons.

The archived BLM and HGO cubes expanded out of plane (see **Figure 6(A2)**). At axial stretch 1.50, the updated CFN cubes contracted in both transverse directions and lost volume in both fibre orientations (see **Figure 6(B)**). The nearly incompressible GOH cube showed little net volume change.

The section-average axial stress was calculated by dividing the axial force by the deformed cross-sectional area of the gauge section (see **Appendix D**). For aligned transverse stretches, the corresponding area relation is:

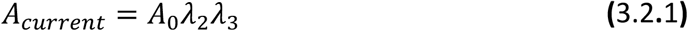

The area-based stress was compared with the estimate obtained by assuming incompressibility:

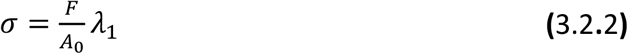

which assumes a constant specimen volume [3, 48]. **Figure 7(A1)** shows the axial stress calculated using the deformed cross-sectional area. Equations **(3.2.1)** and **(3.2.2)** give the deformed-area relation and the stress estimate based on incompressibility, respectively.

**Figure 7.**
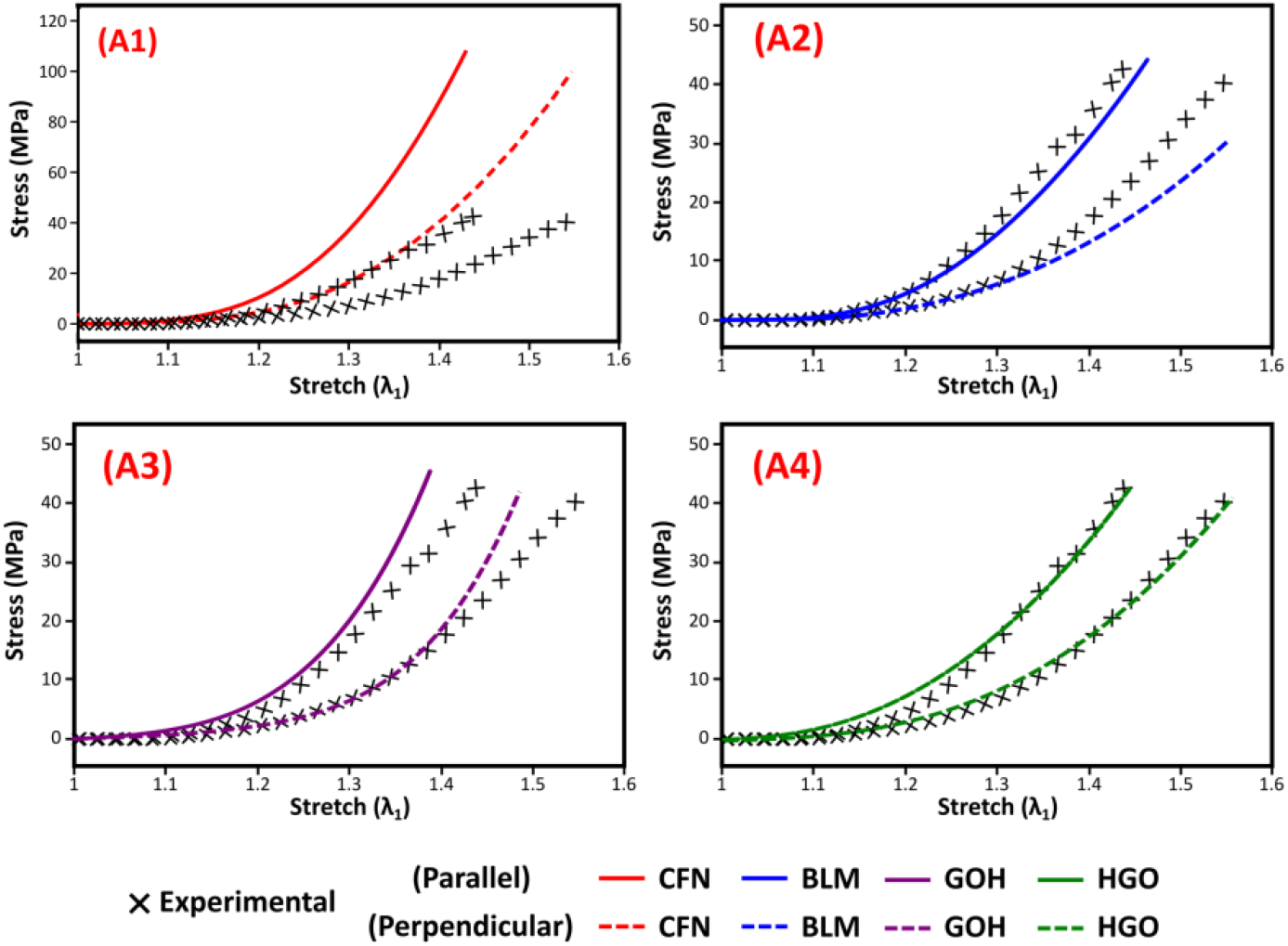
Axial stress–stretch responses. **(A1)** CFN stress calculated using the deformed cross-sectional area over the measured stretch intervals. This panel uses a higher stress-axis limit. **(A2)** BLM, **(A3)** GOH and **(A4)** HGO show the earlier comparison curves. Crosses mark the experimental data. Solid and dashed lines indicate parallel and perpendicular fibre orientations.

### 3.3 Constitutive Model Response Across Different Loading Modes

**Figure 9** shows the indentation fields from the earlier simulations. With the updated CFN parameters, volume ratios ranged from 0.541 to 1.012 and pressures from −0.44 to 2.40 MPa at the prescribed 5 mm displacement, with positive pressure indicating compression.

**Figure 8.**
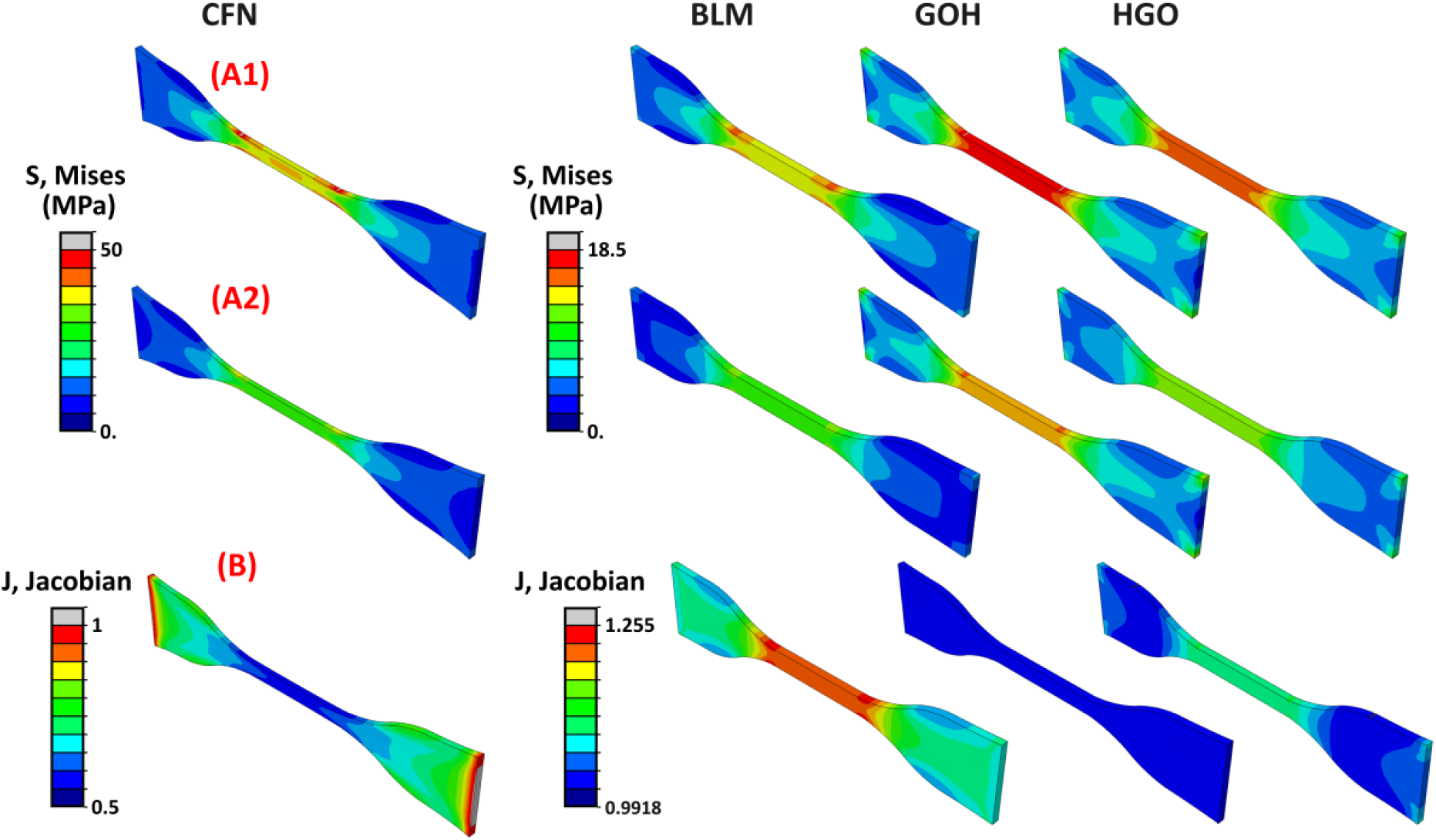
Von Mises stress and volume ratio in the dogbone specimens under uniaxial tension. **(A1)** Parallel and **(A2)** perpendicular von Mises stress. **(B)** Volume ratio for the parallel case. The parallel and perpendicular contours show 40% and 55% elongation, respectively. The CFN contours are from simulations before recalibration.

**Figure 9.**
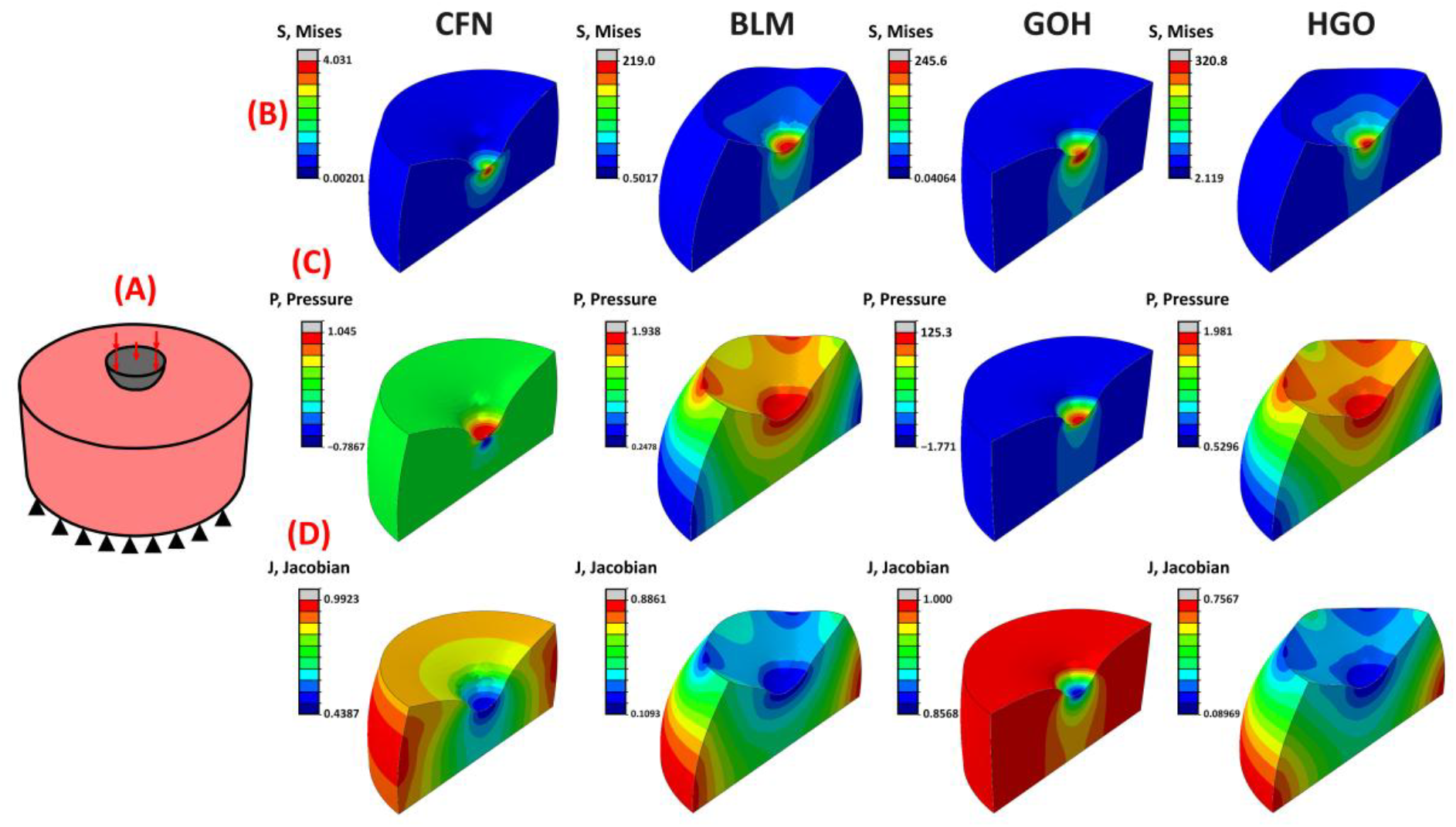
Indentation of the cylindrical specimen in the earlier comparison runs. **(A)** Setup with a 5 mm target displacement. **(B)** Von Mises stress (MPa), **(C)** hydrostatic pressure (MPa) and **(D)** volume ratio. The GOH contours show the response before the target displacement was reached.

The nearly incompressible GOH simulation reached 2.99 mm of indentation, about 60% of the target displacement. The different indentation depths and colour scales limit direct comparison of the contour values.

Additional loading cases, including simple shear and biaxial tension, are presented in **Appendix E**.

## 4 Discussion

The CFN model provides a continuum framework for representing the anisotropic and compressible behaviour of skin. It combines a three-dimensional distribution of collagen fibre orientations with a weak neo-Hookean ground matrix and a separate volumetric potential. The ground matrix has a prescribed shear modulus of 0.1 MPa and provides shear resistance when the fibres are inactive, as shown by the simple-shear example in Appendix F. The volumetric potential describes the tissue’s resistance to changes in volume [49-52]. Volume change also affects fibre stretch because the stretch is calculated from the full deformation. Averaging the response over the reference fibre directions links this directional reinforcement to the continuum stress [53].

The CFN volumetric response was calibrated against the mean uniaxial volume-change curve digitised from Wahlsten et al. [28]. At 20% axial elongation, the parallel CFN simulation gave a volume ratio of 0.711, compared with 0.697 from the published curve (**Figure 3(C2)**). The directional tensile response in **Figure 3(C1)** was fitted to data from a separate study. At larger stretches, the extrapolated volume change depends on the form of the volumetric law. Orientation-dependent volume changes reported elsewhere [54] motivate experiments that measure force, lateral stretch and volume in the same specimens.

Two-family formulations represent preferred directions through prescribed fibre families [35]. The CFN distributes reinforcement over three-dimensional directions informed by reported collagen architecture [20, 22]. The fibre orientation and recruitment parameters were fitted to the tensile curves in both directions (**Figure 3**). The same parameter set described both tensile responses over the calibration range. The fitted distribution had an out-of-plane to in-plane weighted-variance ratio of 29.8%, a statistic calculated from the modelled fibre directions. Under tension, the fibre law represents recruitment through a transition from low to high tensile stiffness [55]. As shown in **Figure B1**, changing the fibre stiffnesses and transition strains alters the initial compliance and subsequent stiffening [55]. In the angular sweep at a stretch of 1.50 (**Figure 5**), the CFN and BLM cubes reached their peak von Mises stress along the prescribed preferred direction. As the loading angle changes, fibres at intermediate orientations can enter or leave tension. A continuous orientation distribution may therefore help describe how stress varies with loading angle. Each curve was normalised by its own peak stress, so assessing accuracy in stress magnitude would require measurements at several loading angles. The simulations in Section **3.3** and **Appendix E** show how the selected models respond under other loading conditions.

The ground matrix contributes shear resistance through its isochoric response, while the separate volumetric potential provides resistance to volume change. This potential builds upon the piecewise framework of Fereidoonnezhad et al. [44]. As shown in **Figure B3**, the intermediate compression branch and the onset of rapid stiffening influence predicted volume change under tension. Beyond the measured range, the terminal branches extend the response according to the selected parameters. The potential contributes isotropic stress [28, 51, 52]. Force, lateral deformation and volume measurements on the same specimens would help distinguish the roles of fibre reinforcement, matrix stiffness and volume change in transverse deformation [41]. At axial stretch 1.50, the CFN cubes contracted in both transverse directions, while the BLM and HGO cubes from the earlier simulations expanded out of plane (**Figure 6(A2)**). A nearly incompressible cube can change shape with little net change in volume (**Figure 6(B)**) [51]. The volume change in **Figure 6(B)** was calculated for the entire cube. In the CFN, volume changes also influence fibre stress through the full fibre stretch [42].

**Figure 7** illustrates how the deformed gauge area affects the calculated axial stress for CFN. At the same axial force and stretch within each simulation, assuming incompressibility overestimated the deformed area and underestimated the section-average axial stress for the contracting CFN specimen. For an expanding specimen with aligned, uniform stretches, the area estimate would shift the calculated stress in the opposite direction. For uniform, aligned stretches, the current area equals the incompressible area multiplied by the volume ratio. At a fixed axial force, the stress calculated using the incompressible area therefore equals the stress calculated using the current area multiplied by the same ratio. The area assumption can thus shift the stress–stretch curve even when the axial force is identical. Using the current gauge area helps distinguish differences in the force carried from differences in the area over which that force acts.

The indentation, shear and biaxial simulations illustrate the response of the selected formulations beyond uniaxial tension. The earlier indentation contours in **Figure 9** show local deformation and the spatial variation in pressure. The GOH contour shows the response before the target displacement was reached (**Figure 9**). Together, **Figure 8** and **Figure 9** illustrate stress and volume distributions in the earlier tension and indentation runs. Fibre recruitment and disengagement can contribute to tension–compression asymmetry [56-58].

The earlier shear comparison in **Figure E1** shows where equivalent stress and volume change are concentrated. These equivalent stresses describe the local response, while the axial stress in **Figure 7** is averaged over a cross-section. The simple-shear contours in **Figure E1(B,C)** show the fields at the displacements used in the earlier simulations. Fibre reorientation and recruitment can influence the response to shear [59]. **Figure E2** shows the earlier biaxial comparison, and the volume ratios in **Figure E2(C)** vary across the cruciform specimen. The models differ in their fibre orientation distributions (**Figure 5**) and in their fibre, matrix and volumetric laws. Given the variability of biological tissues, the response across loading modes requires experimental assessment [60].

Discrete fibre network models can represent individual fibres and non-affine motion, at additional computational cost [61]. The CFN represents fibre reinforcement and compressibility within a continuum formulation [49]. Under pure dilation, increasing the number of sampled fibre directions reduced the anisotropy ratio (**Figure C1(A)**). A resolution of 200 fibre directions was selected to balance directional resolution and computational cost. For this discretisation, computation time increased with the number of elements (**Figure C1(C)**). These results describe the effect of directional sampling and mesh size on the reported benchmark. Spatial mesh convergence requires a separate assessment.

The CFN model is rate independent and describes the ground matrix through an effective isotropic modulus. The unimodal orientation distribution also limits the representation of multiple fibre populations observed in histological studies [20]. The fitted directional parameters describe effective reinforcement. Different parameter sets may reproduce similar calibration curves, and comparison of fit quality across formulations requires equivalent optimisation. Damage and permanent deformation are outside the present formulation and become relevant when modelling tissue failure [62-65]. Independent measurements of lateral deformation are needed to assess the transverse predictions. The stress and volume fields correspond to the simulation increments shown. The preliminary mesh assessment and numerical warnings limit their interpretation.

Experimental assessment should begin by measuring force, axial and lateral deformation, and volume on the same specimens under a common loading protocol. Indentation force– displacement measurements and matched shear or biaxial tests would assess the response under other loading conditions. Varying the volumetric law while holding the fibre and matrix terms fixed would help identify its contribution to the predicted deformation. Imaging the same specimens could constrain the fibre orientation distribution independently of the tensile response.

Extensions to viscoelasticity or poroelasticity would allow time-dependent responses to be investigated [66, 67]. Longer-term processes such as tissue growth and healing would require further modelling and experimental evidence [68, 69].

## 5 Conclusion

The CFN formulation provides a continuum approach for combining orientation-weighted fibre reinforcement, a phenomenological volumetric potential and a weak neo-Hookean ground matrix with a prescribed shear modulus of 0.1 MPa. Calibration used published uniaxial tensile and volume-change data. The updated calibration was applied to indentation, simple shear and biaxial loading. Its piecewise functions provided separate control of fibre stiffening and volumetric resistance. Transfer across anatomical regions and loading conditions requires further assessment.

## Acknowledgment

This project has received funding from the European Union’s Horizon Europe research and innovation programme under grant agreement No 101047008 (BIOMET4D). Views and opinions expressed are however those of the author(s) only and do not necessarily reflect those of the European Union or the European Innovation Council and SMEs Executive Agency (EISMEA). Neither the European Union nor the EISMEA can be held responsible for them.

## CRediT Authorship Contribution Statement

Conall Quinn: Conceptualisation, Methodology, Software, Formal Analysis, Investigation, Visualisation, Writing – original draft.

Mahtab Vafaeefar: Writing – review & editing.

Behrooz Fereidoonnezhad: Writing – review & editing.

Ted J. Vaughan: Supervision, Funding Acquisition, Project Administration, Writing – review & editing.

## Data availability

The experimental data were obtained from the cited publications. Model inputs and supporting scripts are available from the corresponding author on reasonable request.

## Appendix A CFN Volumetric Formulation

### A.1 Compression (J < 1)

Compression used linear, quadratic, nonlinear and terminal exponential branches. The thresholds *J*_1*c*_, *J*_2*c*_ and *J*_3*c*_ satisfied 0 < *J*_3*c*_ < *J*_2*c*_ < *J*_1*c*_ = 1. The linear branch applied over *J*_1*c*_ ≤ *J* ≤ 1, and the quadratic transition over *J*_2*c*_ ≤ *J* < *J*_1*c*_:

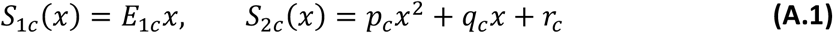

Here *x* = *J* − 1 and *x*_*jb*_ = *J*_*jb*_ − 1 for *j* ∈ {1,2,3}, *b* ∈ {*c, e*}. Matching the value and first derivative gave:

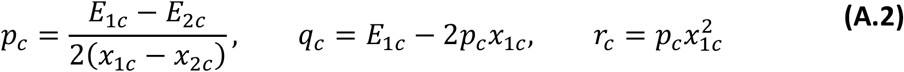

For *J*_3*c*_ ≤ *J* < *J*_2*c*_, set *y*_*c*_ = *x* − *x*_2*c*_ and *B*_2*c*_ = *S*_2*c*_(*x*_2*c*_). The fixed stress reference *S*^*^ = 1 MPa made the logarithm dimensionless. The nonlinear branch was:

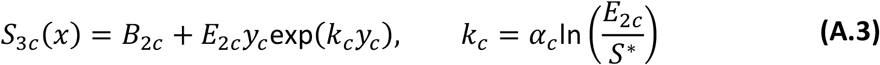

For *J* < *J*_3*c*_, set *z*_*c*_ = *x* − *x*_3*c*_ and *B*_3*c*_ = *S*_3*c*_(*x*_3*c*_). The terminal branch was:

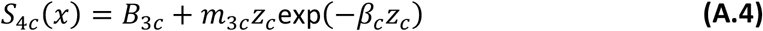

With *y*_3*c*_ = *x*_3*c*_ − *x*_2*c*_, the matched slope was:

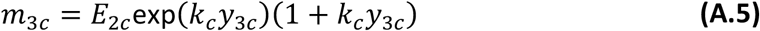

### A.2 Expansion (J > 1)

Expansion used the same four-branch structure. Its thresholds *J*_1*e*_, *J*_2*e*_ and *J*_3*e*_ satisfied 1 < *J*_1*e*_ < *J*_2*e*_ < *J*_3*e*_. The linear branch applied over 1 < *J* ≤ *J*_1*e*_, followed by the quadratic transition over *J*_1*e*_ < *J* ≤ *J*_2*e*_:

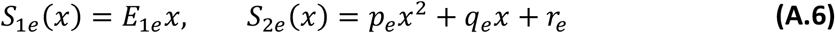

With *x* = *J* − 1, the matching coefficients were:

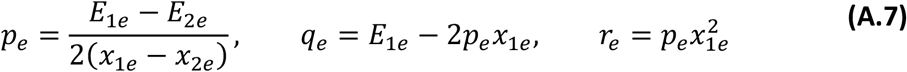

For *J*_2*e*_ < *J* ≤ *J*_3*e*_, set *y*_*e*_ = *x* − *x*_2*e*_ and *B*_2*e*_ = *S*_2*e*_(*x*_2*e*_). The tabulated value *α*_*e*_ = 0 gave:

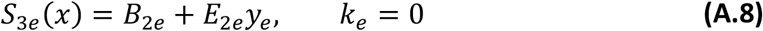

For *J* > *J*_3*e*_, set *z*_*e*_ = *x* − *x*_3*e*_ and *B*_3*e*_ = *S*_3*e*_(*x*_3*e*_). The terminal exponential branch was:

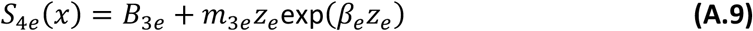

The terminal-branch slope was:

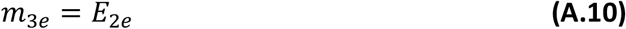

Stress and its first derivative matched at the internal branch joins. The compression and expansion tangents at unit volume ratio were distinct. **Figure A1** shows the expansion law evaluated with the coefficients in Table 3.1.1.

**Figure A1.**
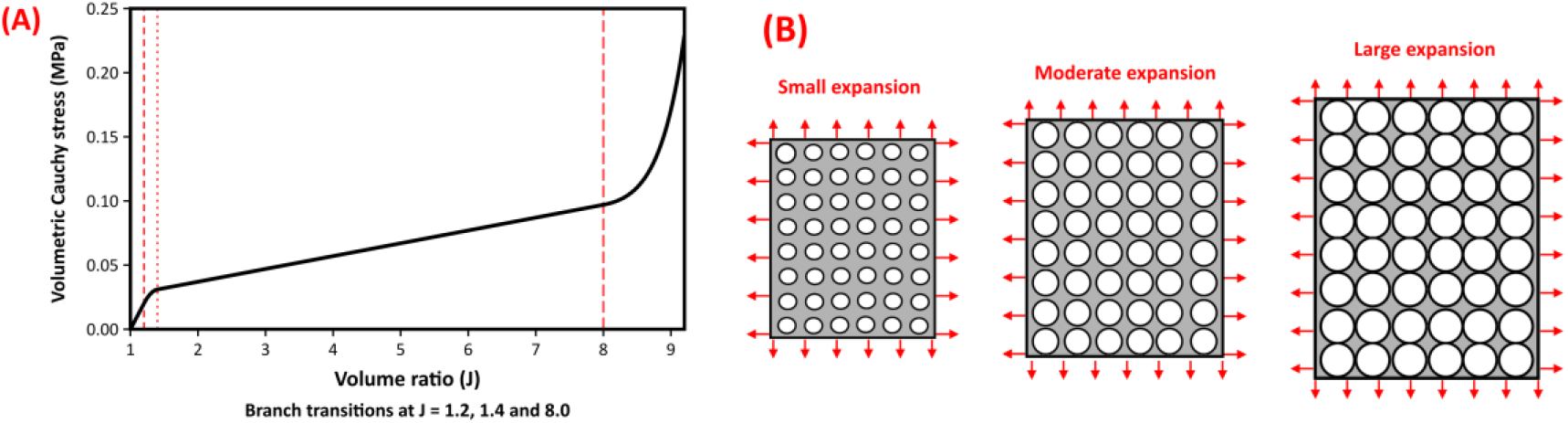
Volumetric stress response under expansion (*J* > 1). (A) Volumetric Cauchy stress from the prescribed expansion law. Dashed guides mark the branch transitions. (B) Schematic illustration of increasing volume.

## Appendix B CFN Parameter Sensitivity Analysis

Sensitivity analyses were performed on fibre mechanical, architectural, and volumetric parameters using a unit cube under uniaxial tension. Each sweep varied the indicated parameter or orientation axes and held the other settings at the joint calibration in **Table 3.1.1**.

### B.1 Fibre Mechanical Parameters

The impact of collagen fibre mechanics was evaluated by varying uncoiling stiffness *E*_1*f*_, first transition strain *D*_1*f*_, straightening stiffness *E*_2*f*_, and second transition strain *D*_2*f*_. As shown in **Figure B1** variations in *E*_1*f*_ primarily affected the initial compliant region of the stress– stretch response, with higher values increasing stiffness at low stretches (**Figure B1(A)**). Changes in *D*_1*f*_ shifted the transition from uncoiling to stiffening, thereby delaying or advancing fibre recruitment (**Figure B1(B)**). The parameter *E*_2*f*_ governed the slope of the stress response at larger stretches (**Figure B1(C)**), while *D*_2*f*_ controlled the onset of the fully straightened fibre regime (**Figure B1(D)**). The difference between parallel and perpendicular responses reflects the prescribed fibre orientation.

**Figure B1.**
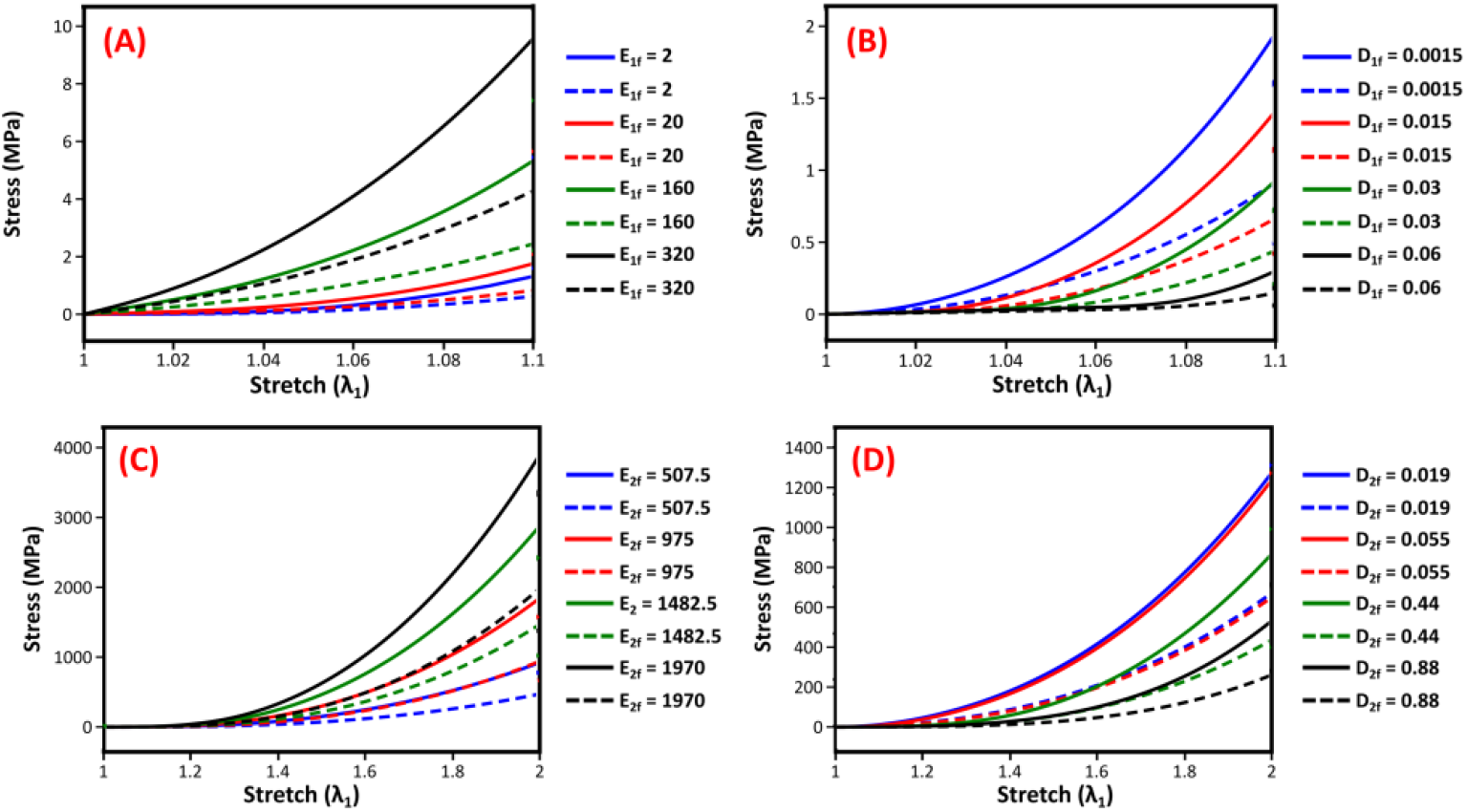
Sensitivity analysis of the CFN model examining the effect of collagen fibre mechanics on the stress-stretch response. **(A)** Uncoiling modulus (*E*_1*f*_) and **(B)** first transition strain (*D*_1*f*_) influence early compliance. **(C)** Straightening modulus (*E*_2*f*_) and **(D)** second transition strain (*D*_2*f*_) govern stiffness at higher stretches. Solid and dashed lines represent parallel and perpendicular fibre orientations, respectively. The coefficients outside each sweep were held at their jointly calibrated values (see **Table 3.1.1**).

### B.2 Fibre Orientation Parameters

The influence of fibre network architecture was evaluated by varying the alignment parameters (*α*_*x*_, *α*_*y*_, *α*_*z*_) under uniaxial tensile loading. The parameter *α*_*x*_ controlled preferential alignment along the loading direction, while *α*_*y*_ and *α*_*z*_ regulated dispersion in the transverse and out-of-plane directions. The orientation weights summed to one in each case. Increasing *α*_*x*_ increased directional bias and tissue stiffness in the loading direction. Reducing *α*_*y*_ and *α*_*z*_ further enhanced anisotropy by limiting transverse and out-of-plane fibre support. A uniform distribution (*α*_*x*_ = *α*_*y*_ = *α*_*z*_) gave a smaller difference between the parallel and perpendicular responses. **Figure B2** shows the responses to these changes in alignment.

**Figure B2.**
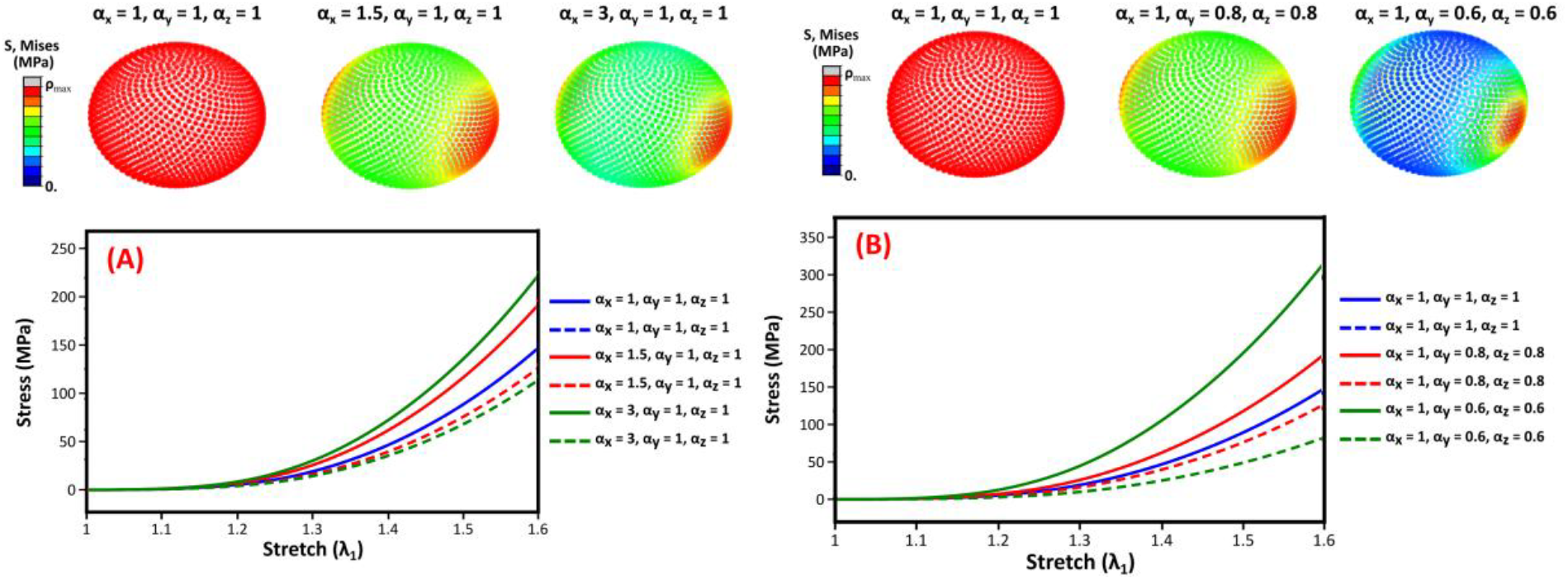
Sensitivity analysis of the CFN model altering the **(A)** preferential fibre direction (*α*_*x*_) and **(B)** non-preferential fibre directions (*α*_*y*_, *α*_*z*_) with collagen fibres orientated in the parallel (solid line) and perpendicular (dashed line) directions. The upper panels show the original orientation illustrations. Other coefficients in the stress curves were held at their fitted values (see **Table 3.1.1**).

### B.3 Sensitivity to Volumetric Parameters

The intermediate-branch coefficient affected the compression law and volume change during tension (**Figure B3(A,B)**). The effect of this coefficient on volume loss varied with stretch. The compression stress curves crossed because the coefficient also enters the nonlinear scaling. Increasing the terminal-branch threshold brought forward rapid stiffening and reduced volume loss at larger tensile stretches (**Figure B3(C,D)**).

**Figure B3.**
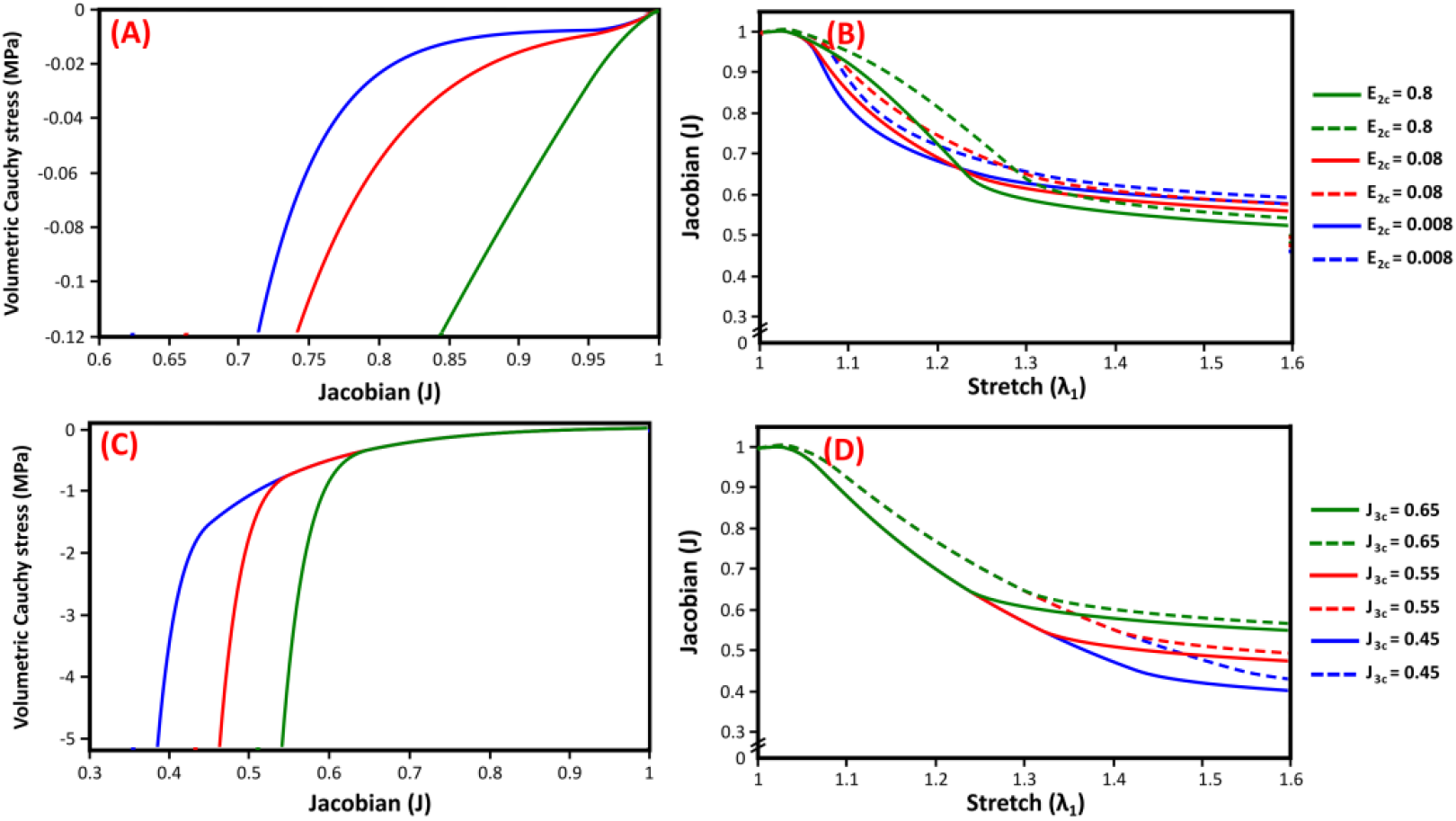
Volumetric sensitivity of the updated CFN model. **(A)** Prescribed compression stress and **(B)** cube volume ratio for the intermediate-branch coefficient sweep. **(C)** Prescribed compression stress and **(D)** cube volume ratio for the terminal-branch threshold sweep. Coefficients outside each sweep were held at the values in **Table 3.1.1**.

## Appendix C Fibre Number Sensitivity and Computational Cost

**Figure C1** reports the original fibre-count and computational-cost benchmark, which used its own simulation settings. Increasing the number of fibres reduced the anisotropy ratio (χ) under pure dilation, indicating diminished directional bias associated with discrete fibre sampling. The ratio decreased approximately linearly on a logarithmic scale between 20 and 500 fibres. An isotropy threshold of **χ** < 0.01 was adopted as a criterion for negligible directional bias. Higher counts further reduced χ below the stated threshold and increased computational cost. Networks with 20, 100, and 500 fibres maintained stable CPU times of 137.3, 136.2, and 146.4 seconds respectively, whereas fibre counts of 1000 and 10,000 resulted in a marked increase in computational time (**Figure C1(B)**). All simulations were executed on a workstation equipped with an AMD Ryzen Threadripper 3960X processor (24 cores, 48 threads, 3.79 GHz).

CFN exhibited CPU times comparable to the BLM (139.2 s), GOH (134.7 s), and HGO (140.1 s) models, despite representing up to 500 discrete fibre directions compared with 36, 2, and 2 directions in those formulations. Fibre counts between 100 and 500 satisfied the isotropy criterion while preserving computational efficiency. The updated CFN cases retained the 200-direction discretisation. **Figure C1(C)** shows the increase in CPU time with element number for the 200-fibre configuration.

**Figure C1.**
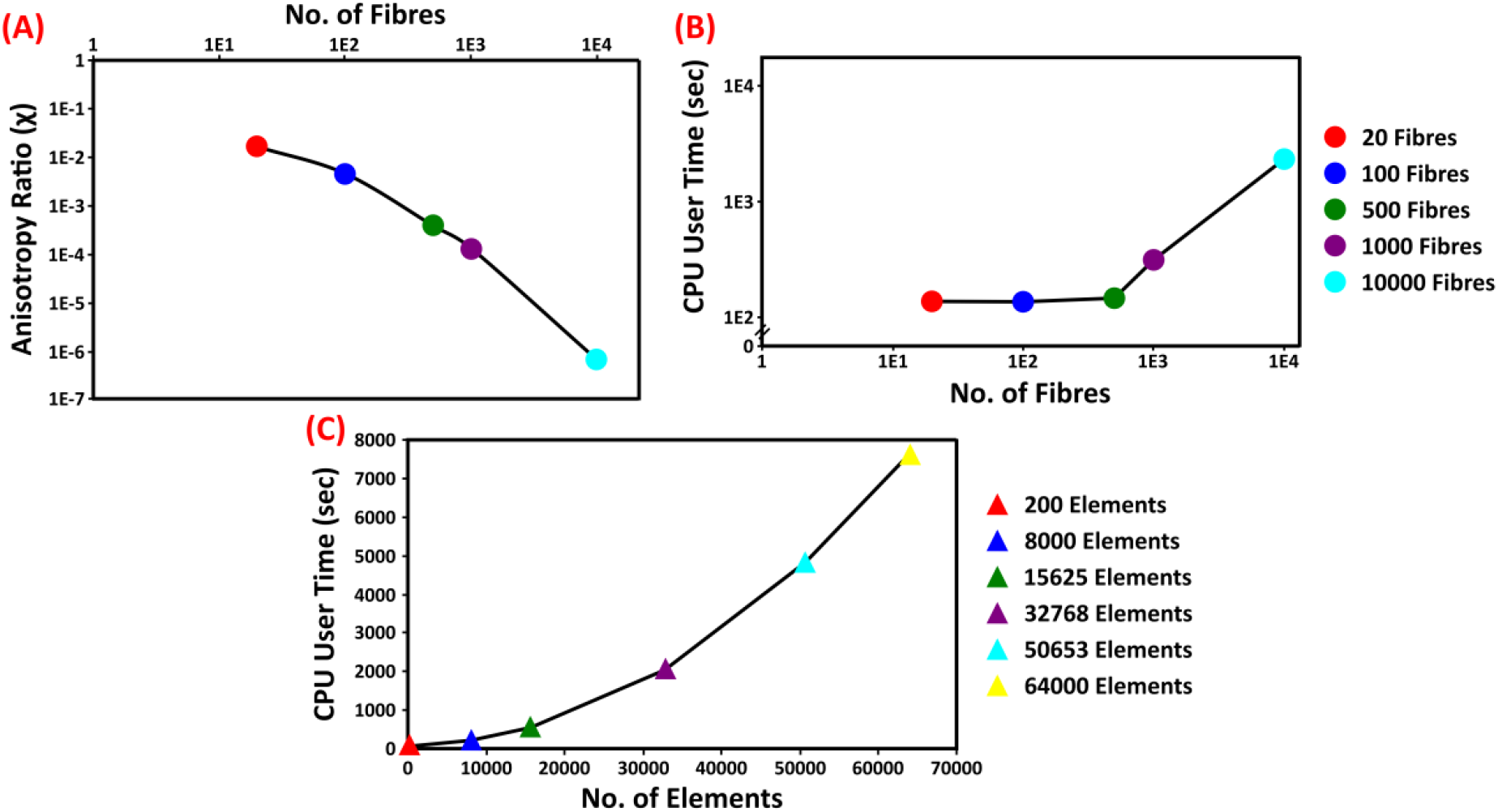
Impact of fibre number on **(A)** anisotropy ratio [1.63*E* − 2, 4.57*E* − 3, 3.86*E* − 4, 1.33*E* − 4, 6.718*E* − 07] and **(B)** CPU user time [137.3, 136.2, 146.4, 313.4, 2328.4] at 20, 100, 500, 1000 and 10000 fibres respectively. **(C)** Effect of increasing element number on user CPU time at 200 fibres at 200, 8000, 15625, 32768, 50653 and 64000 elements.

## Appendix D Instantaneous Cross-Sectional Area Computation

The instantaneous area of the gauge section was computed from the deformed finite element geometry to estimate the section-average axial stress.

Nodal coordinates defining the specimen cross-section were extracted at each increment. The four nodes were taken in perimeter order. The quadrilateral was split along the diagonal from the first to the third node, and the two triangle areas were summed:

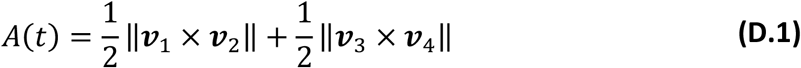

where ***v***_1_, ***v***_2_, ***v***_3_, ***v***_4_ are edge vectors formed from the nodal positions (***x***_1_, ***x***_2_, ***x***_3_, ***x***_4_) as:

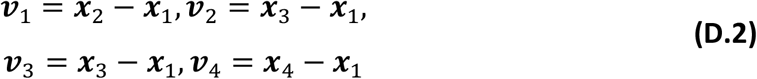

Dividing the axial force by this area accounts for the evolving gauge geometry without assuming incompressibility. **Figure D1** shows the tracked nodes and the evolution of the gauge area.

**Figure D1.**
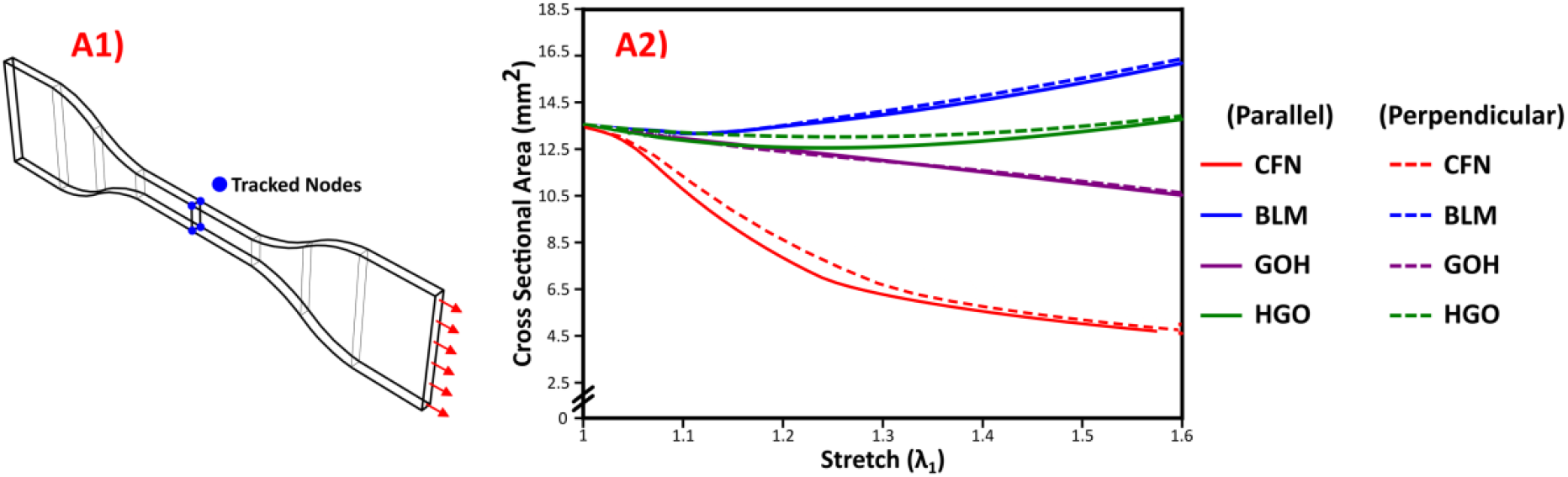
**(A1) Visualisation** of tracked nodal positions used to compute instantaneous cross-sectional area. **(A2) Evolution** of cross-sectional area under uniaxial tension for the updated parallel and perpendicular CFN dogbones, plotted against measured gauge stretch alongside the archived comparison curves.

## Appendix E Multi-Axial Comparative Model Behaviour

**Figure E1(B)** compares the equivalent stress fields under simple shear. CFN and GOH developed localised stress concentrations, while BLM and HGO distributed stress more broadly through the specimen. The volume fields in **Figure E1(C)** show contraction across much of the CFN specimen, near-constant volume for GOH, and regions of contraction and expansion for BLM and HGO. The peak stresses reflect different prescribed displacements.

In the separate 5 mm simulation using the joint CFN calibration, the integration-point von Mises stress ranged from 0.021 to 45.49 MPa and the volume ratio ranged from 0.626 to 1.005.

**Figure E1.**
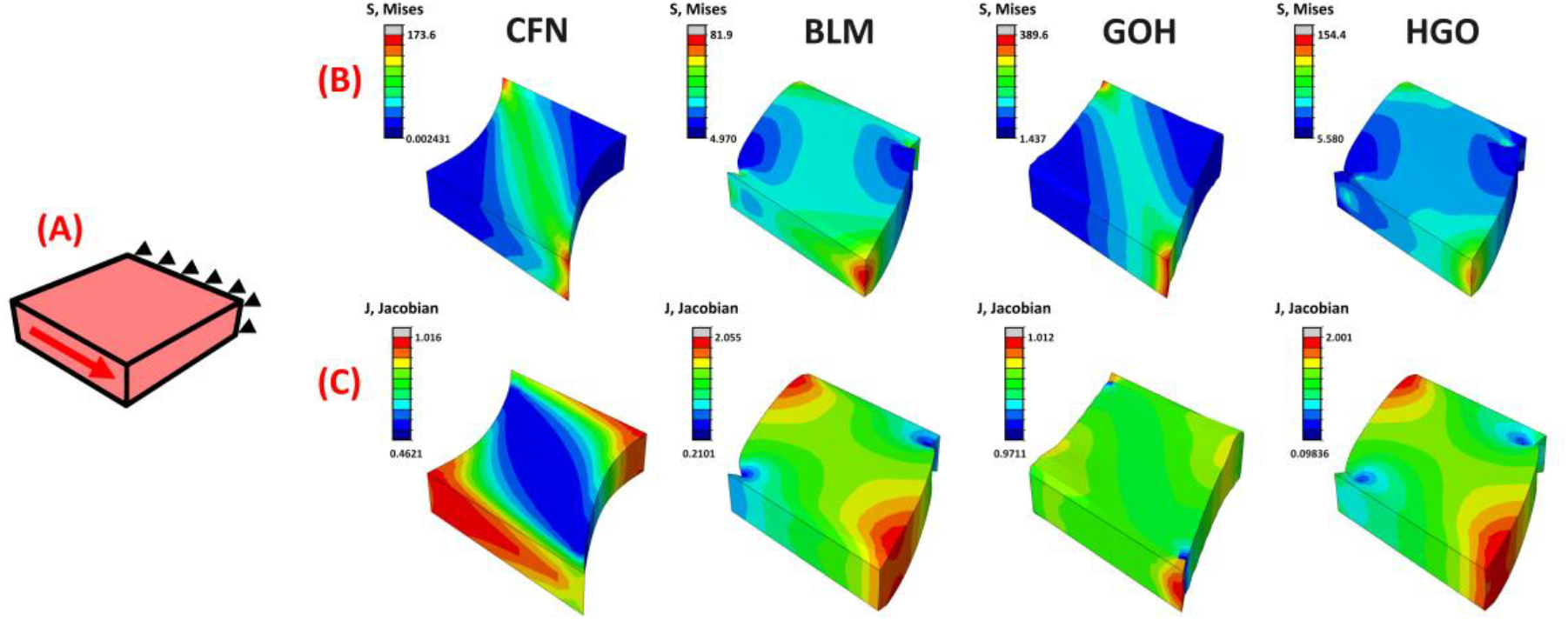
Earlier simple-shear comparison. **(A)** Loading schematic. **(B)** Von Mises stress (MPa). **(C)** Volume ratio. The contours show the earlier simulations at their respective displacements. Limits describe the displayed fields.

The models produced different stress patterns under biaxial loading (**Figure E2(B)**). The CFN contour showed volume loss across much of the cruciform specimen, with local expansion near its boundaries. GOH remained close to constant volume, while the displayed BLM and HGO fields showed expansion. In the separate biaxial simulation using the joint CFN calibration, the volume ratio ranged from 0.653 to 1.159, with contraction in most elements and local expansion.

**Figure E2.**
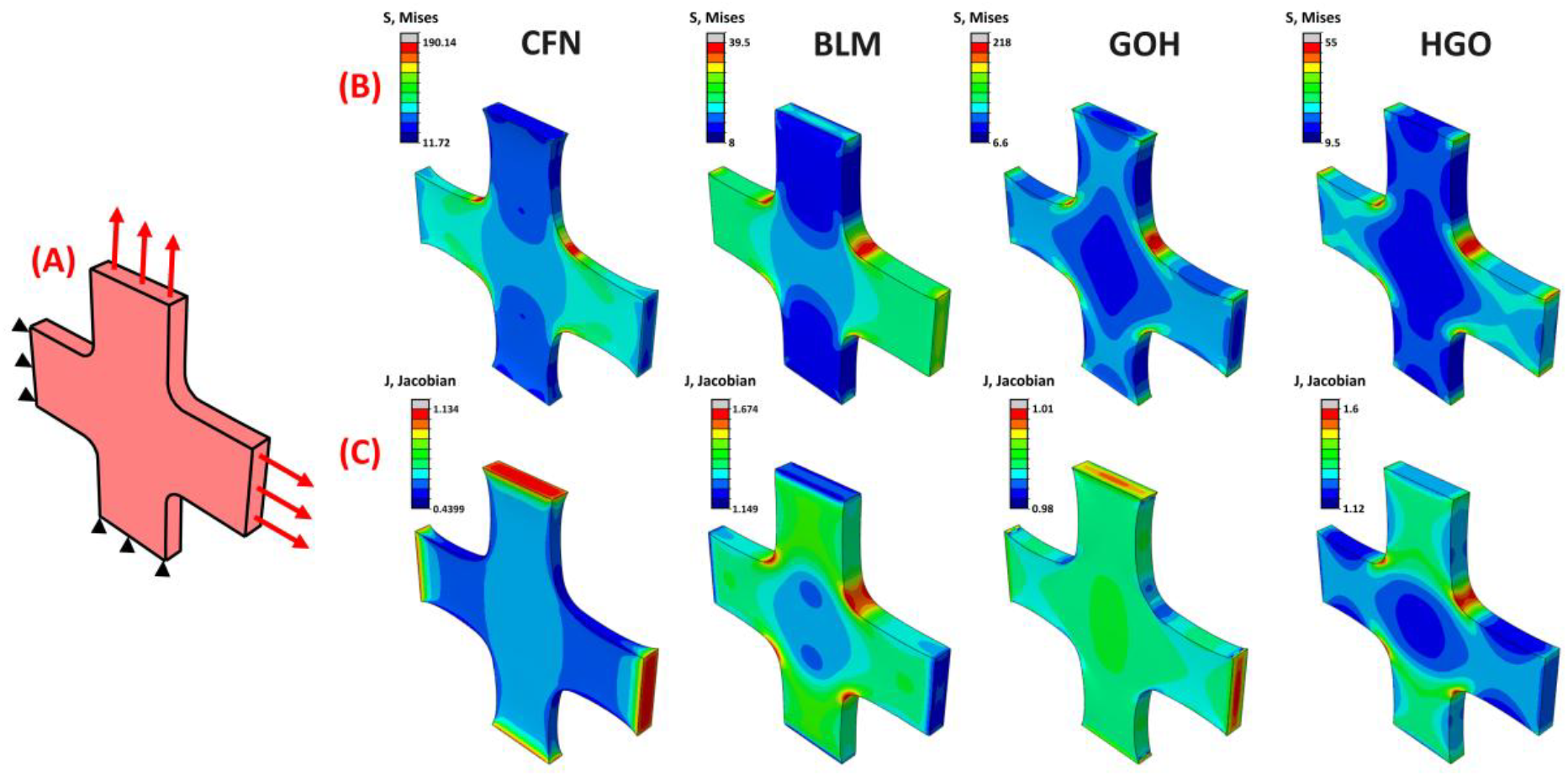
**(A)** Earlier biaxial comparison, with opposing grips displaced by 2.5 mm per edge along each axis. **(B)** Von Mises stress (MPa) with *σ*_*max*_ = [190.14, 39.5, 218, 55] and *σ*_*min*_ = [11.72, 8, 6.6, 9.5]. **(C)** Jacobian *J*_*min*_ = [0.4399, 1.149, 0.98, 1.12] and *J*_*max*_ = [1.134, 1.674, 1.01, 1.60]. These display limits are for CFN, BLM, GOH and HGO, respectively.

## Appendix F Strain Energy Derivations

### F.1 Energy and stress measures

Differentiate the reference-volume energy in Equation **(2.2.1)** at fixed fibre directions and orientation probabilities to obtain the second Piola–Kirchhoff and Cauchy stresses.

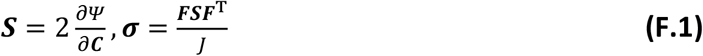

### F.2 Fibre energy

Integrating the fibre coefficient in Equation **(2.2.16)** gives the fibre energy.

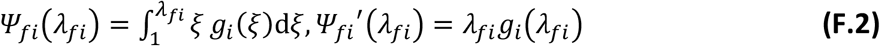

The energy vanishes at and below unit stretch. Using 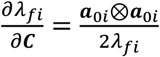 in (F.1) gives the fibre-network stress in Equation **(2.2.18)** with one factor of *p*_*i*_ per direction.

### F.3 Volumetric and matrix energies

Integrating the signed stress coefficient from Equation **(2.2.19)** and Appendix A gives the volumetric energy.

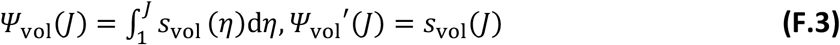

Using 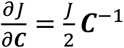 in (F.1) gives *s*_vol_(*J*)***I***. The matrix energy in Equation **(2.2.20)** gives the deviatoric stress.

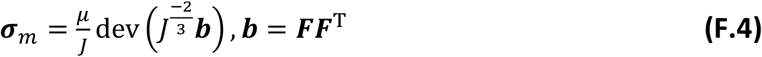

Adding the three contributions gives the combined Cauchy stress.

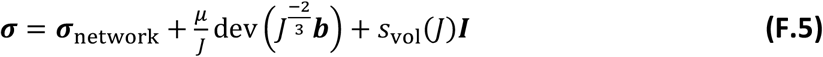

Both integrals carry the accumulated energy across branch joins. Energy and its first two derivatives match at internal joins. At fibre activation and *J* = 1, the first derivative is continuous and the second changes. The reference energy is zero.

For isochoric simple shear ***F*** = ***I*** + *γ****e***_1_ ⊗ ***e***_2_, a fibre aligned with ***e***_1_ stays at unit stretch. The matrix contributes energy ^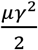^ and shear stress ***σ***_*m*,12_ = *μγ*.

